# Decoding Amyloid Plaque Penetrability: Exploring Extracellular Space and Rheology in Plaque-rich Cortex

**DOI:** 10.1101/2025.08.24.671960

**Authors:** Juan Estaún-Panzano, Yulia Dembitskaya, Ivo Calaresu, Somen Nandi, Quentin Gresil, Evelyne Doudnikoff, Thierry Leste-Laserre, Thierry Amédée, Laurent Cognet, Laurent Groc, U. Valentin Nägerl, Erwan Bezard

## Abstract

A hallmark of Alzheimer’s disease (AD) is the accumulation of amyloid plaques, primarily composed of misfolded amyloid β (Aβ) peptides. We employed complementary high-resolution imaging techniques to investigate the plaque penetrability and the extracellular space (ECS) rheology in a mouse model of AD. Two-photon shadow imaging *in vivo* confirmed that a dense ring of cells surrounds cortical amyloid plaques but highlighted the diffusional penetrability of the amyloid core. Quantum dot tracking unveiled that ECS diffusional parameters are heterogeneous in and around plaques, with an elevated diffusivity within and around plaques compared to WT-tissue. The amyloid core showed low nanoparticle density, varying by plaque phenotype. Carbon nanotube tracking confirmed these altered local rheological properties at the level of the whole cortex of AD mice. Finally, we found the extracellular matrix to be dysregulated within the amyloid plaque, which may account for the observed alterations in diffusivity. Our study provides fresh insights for understanding Aβ plaque penetration, a prerequisite for therapeutic development.

## 1. Introduction

Alzheimer’s disease (AD) poses significant challenges to researchers and clinicians due to its poorly understood aetiology and pathophysiology. This condition profoundly affects cognitive function, especially in the expanding ageing population, becoming a significant socio-economic burden for healthcare systems [1]. A recognised hallmark of the disease is the presence of amyloid plaques, primarily composed of misfolded amyloid β (Aβ) peptides, in the brain’s extracellular space (ECS) [2].

Among recent strategies to tackle the disease, amyloid plaque clearance and immune system modulation have raised hopes but still suffer from unsatisfactory outcomes [3]. As a consequence, the penetrability of brain tissue has come into focus as a potentially pivotal determinant of therapeutic efficacy [1]. However, how different substances can navigate within the ECS remains poorly understood, let alone whether they can penetrate amyloid plaques. Deeper insights into the properties of the ECS are crucial, as it plays a key role in normal brain function [4] and disease progression [5].

This study presents an innovative experimental and analytical framework for dissecting the influence of the ECS rheology, i.e. a concept that encompasses diffusionability within the compartment and compartment dimensions, on molecular diffusion in the context of amyloid pathology. We used 2-photon shadow imaging and nanoscopic single-particle tracking (SPT) imaging to investigate the morphology and diffusional properties of the ECS, i.e. its rheology, in the transgenic APP/PS1 mouse model of AD. Our study provides new insights into the factors that influence the penetration of exogenous molecules into Aβ plaques themselves and Aβ plaques-bearing cortex.

## 2. Results

### 2.1 Shadow imaging reveals amyloid plaque organisation and surrounding ECS structure

We focused our study on the sensory and motor cortices. The amyloid pathology burden starts as early as 3 months of age and displays a significant load of plaques in 8-month-old female APP/PS1 mice [6]. To visualise amyloid plaques in their native environment, we used 2-photon shadow imaging (TUSHI) *ex vivo* in acute brain slices and *in vivo* in anesthetised mice, achieving sub-micron resolution and panoptical visualisation of brain tissue architecture [7]. We labelled the ECS using cell membrane-impermeable dyes (Calcein bath applied in acute brain slices, and Alexa Fluor 488 injected intracerebroventricularly *in vivo*, see Methods for details), projecting all luminal structures (cells, blood vessels) as black “shadows” in the images. For better illustration, we inverted the image contrast so that all cellular structures appear white and the ECS black (**Figure 1a**). In acute brain slices, the approach detected amyloid plaques in the cortex (Supplementary Video 1). Their amyloid region was filled with Calcein (black in the inverted images). The *ex vivo* finding was confirmed *in vivo* after intracerebroventricular injection of Alexa 488 Fluor (Supplementary Video 2,3). Notably, amyloid plaques were frequently associated with blood vessels (Supplementary Video 2, 3), which supports the notion of neurovascular alterations in AD pathology [8]. Both dyes appeared uniformly distributed in the centre of the amyloid plaques, but this does not mean that the ECS in this region is empty (it is chock-full with proteins, lipids, organelles and dystrophic neurites with compromised membranes) [9],[10]. Instead, it shows that small organic dyes with hydrodynamic diameters on the order of a nanometre have no trouble penetrating amyloid plaques, which have an internal structure that cannot be resolved by diffraction-limited light microscopy.

**Figure 1.**
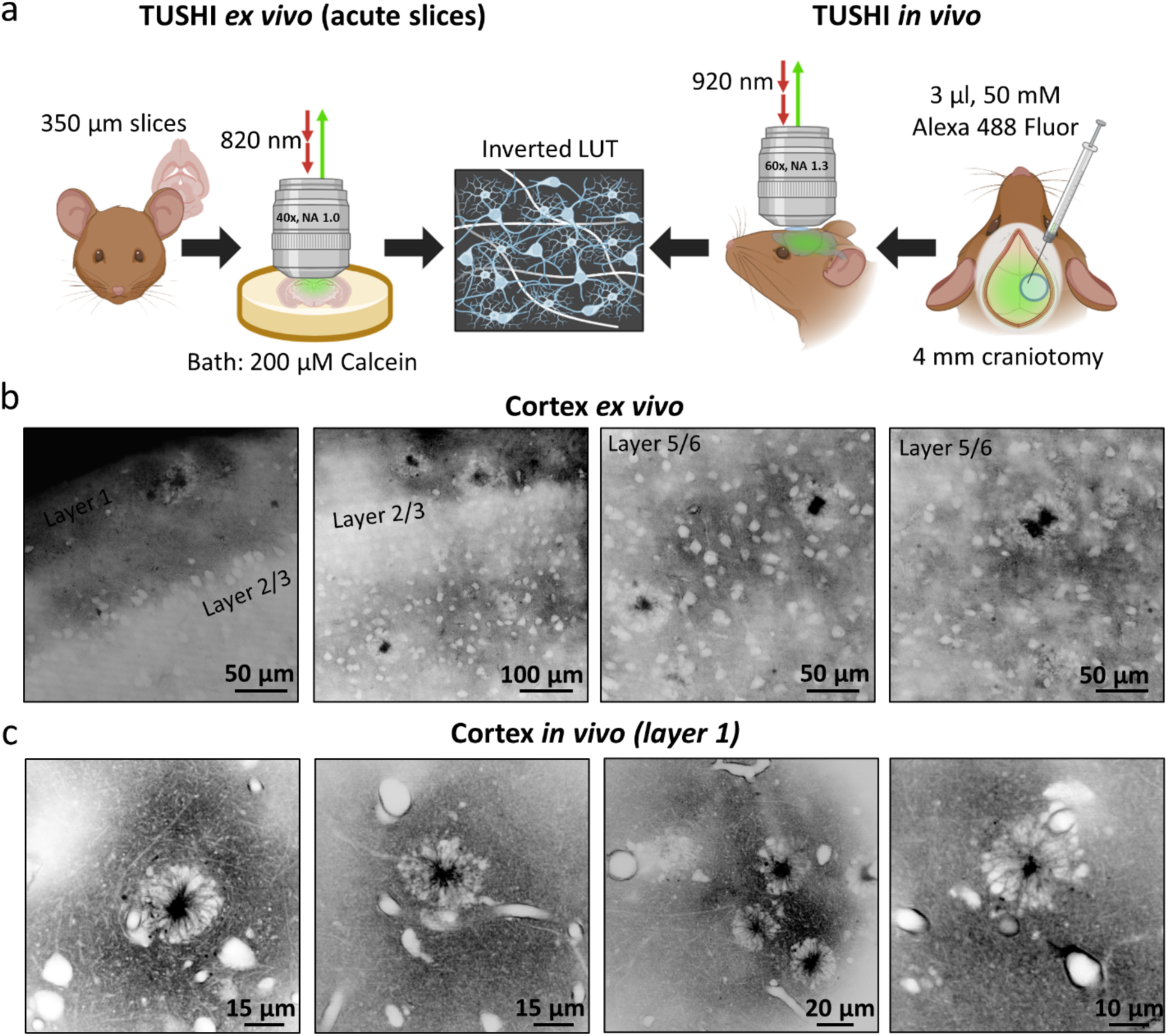
Visualisation of amyloid plaques using TUSHI in cortex *ex vivo* and *in vivo*. a – Schematic representation of visualisation of amyloid plaques using TUSHI in vitro and in vivo. TUSHI *ex vivo*: acute brain slices (350 µm thick) were placed into a microscope chamber after incubation and 200µM of Calcein was bath applied to label extracellular space. TUSHI *in vivo*: Alexa 488 Fluor (3 µl, 50 mM) was injected in the lateral ventricle on the side of 4 mm craniotomy and covered with a glass coverslip afterwards. All collected images are presented in inverted LUT where extracellular space appears black and brain structures - white. b – representative images of amyloid plaques in cortex *ex vivo*. c – representative images of amyloid plaques in the cortex *in vivo*. Created in BioRender. Bezard, E. (2024) BioRender.com/i34k786.

Plaques were irregularly distributed in the brain parenchyma and present in all cell layers of the cortex (layers 1 - 6 (Figure 1b). Notably, they always featured a dense sphere of cells that looked like a ring in optical sections (Supplementary Videos 1-3), surrounding the core of the amyloid plaque, as previously reported in many instances.

We then performed TUSHI through a cranial window in the somatosensory cortex *in vivo* (Figure 1c), which confirmed the ring/amyloid arrangement as a consistent feature in the diseased brain (Supplementary Video 2, 3), which was often associated with blood vessels (Figure 1c). By contrast, we never observed such structures in age-matched wild-type (WT) mice (Supplementary Figure 1, Supplementary Videos 4,5).

We measured the surface area of the amyloid cores and the rings around them *ex vivo* and *in vivo* in the central image section of a plaque (**Figure 2a**). The surface areas of the amyloid cores and cell rings were indistinguishable between the *ex vivo* and *in vivo* conditions (Figure 2b). The surface area of the cell ring was about 6.6 times bigger on average than the surface area of the core (Figure 2c). The surface areas of all measured amyloid cores (Figure 2d) and their cell rings (Figure 2e) follow a log-normal distribution, consistent with X-ray phase-based virtual histology [11].

**Figure 2.**
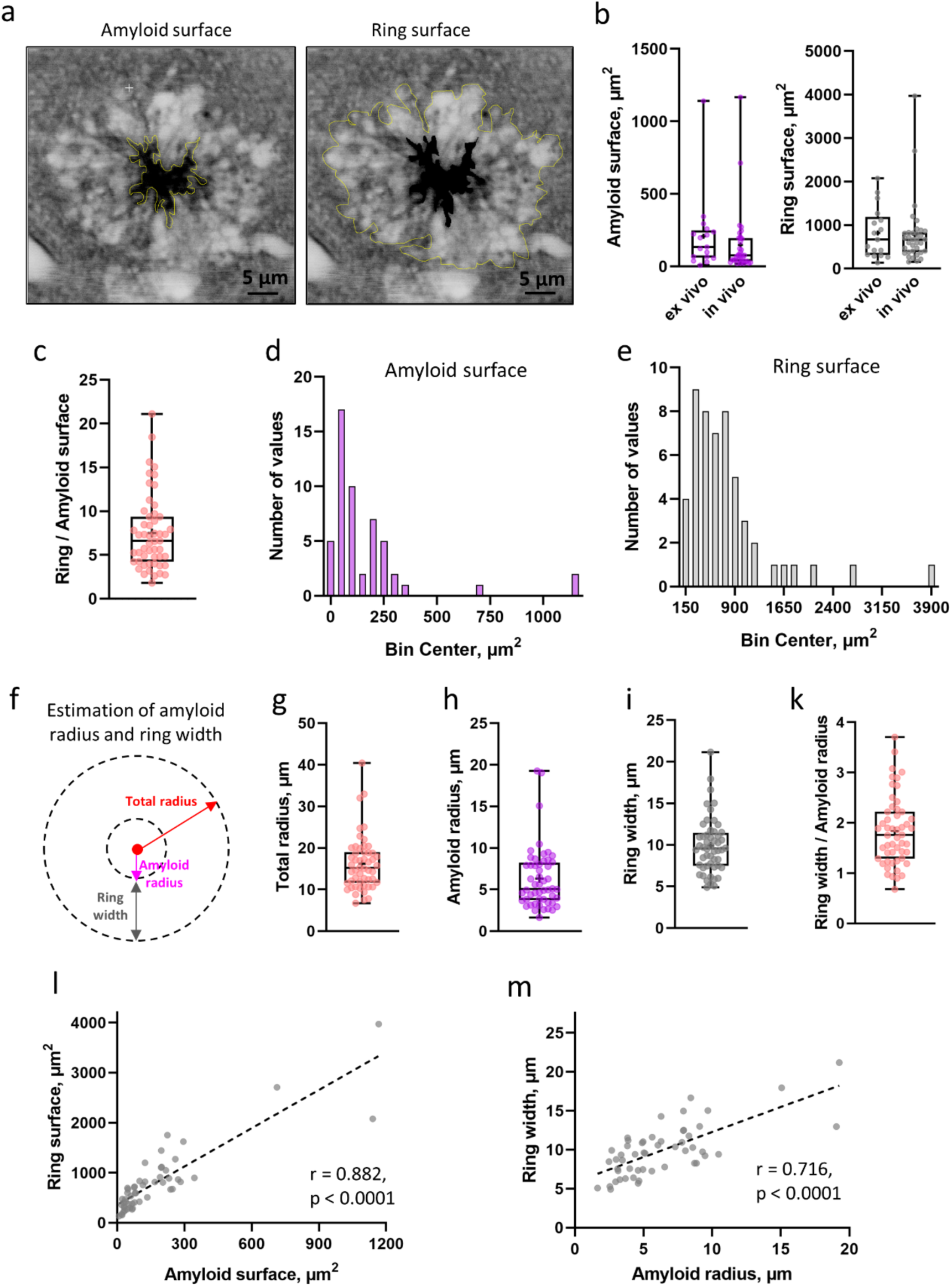
Dimensions of amyloid plaques using TUSHI. a – Manually selected ROIs were calculated using the “free-shape” drawing tool in ImageJ, from which the amyloid and ring surface area were calculated. b - Left panel - Amyloid surface in cortex ex vivo (134.2 µm2, IQR = 61.63 – 248.7, n = 17) and in vivo (77.0 µm2, IQR = 35.6 – 195.6, n = 35) (Mann-Whitney test, p = 0.134), Right panel - Ring surface in cortex ex vivo (666.7 µm2, IQR = 312.3 - 1193, n = 17) and in vivo (666.4 µm2, IQR = 381.5 – 833.1, n = 35) (Mann-Whitney test, p = 0.671). c - The surface of the ring was 6.6 times higher than that of amyloid plaque in the cortex (ex vivo & in vivo) (IQR = 4.24 – 9.36, n = 52). d - log-normal distribution of amyloid surface in the cortex (ex vivo & in vivo) (D’Agostino & Pearson test, p = 0.765, n=52). b – log-normal distribution of ring surface area in the cortex (ex vivo & in vivo) (D’Agostino & Pearson test, p = 0.910, n=52). f - Approximation of amyloid plaques as a circle to calculate amyloid radius and width of the ring. The total radius (amyloid + ring) was calculated from the surface of the amyloid + ring. The width of the ring was calculated as the difference between the total radius and the radius of the amyloid. g – Estimation of total radius of amyloid + ring in cortex (ex vivo & in vivo) (15.21 µm, IQR = 11.62 – 18.98, n= 52). h – Estimated amyloid radius in cortex (ex vivo & in vivo) (5.1 µm, IQR = 3.71 – 8.25, n = 52). i – Estimated width of the ring in the cortex (ex vivo & in vivo) (9.57 µm, IQR = 7.47 – 11.44, n = 52). k – The width of the ring was 1.82 times higher than that of the amyloid radius in the cortex (ex vivo & in vivo) (IQR = 1.29 – 2.22, n = 52). l - Positive correlation of ring and amyloid surface (Pearson’s r = 0.882, p < 0.0001). m - Positive correlation of ring width and amyloid radius (Pearson’s r = 0.716, p < 0.0001). All data are represented as median and IQR, minimum, maximum and mean (labelled as a cross) with individual data points.

Approximating a spherical geometry for the plaques comprised of amyloid core and cell ring, we calculated their radius and width, respectively (Figure 2f). The amyloid radius was 5.1 µm (Figure 2h), while the ring width was 9.57 µm (Figure 2i), making the width 1.82 times bigger than the radius (Figure 2k). The median radius of amyloid plaques (amyloid core + cell ring) was 15.2 µm (Figure 1g). The radius of the amyloid core and the width of the ring were indistinguishable between the *ex vivo* and *in vivo* conditions (Supplementary Figure 2). Lastly, we observed a positive correlation between the surface areas of the amyloid core and cell ring (Figure 2l). Likewise, radius and width were positively correlated (Figure 2m), meaning larger amyloid cores had wider cell rings. Remarkably, these numbers and geometrical assumptions indicate that, on average, the entire plaque structure takes up about 26 times more volume than the amyloid core by itself.

The strength of our novel approach lies in clearly distinguishing between membrane-compromised structures (which the dye penetrates) and intact ones, while also providing precise size measurements of the different regions of the plaque (core *versus* ring). While TUSHI does not provide diffusional information, it allows us to identify regions of interest (ROIs) for further blind quantum dot single-particle tracking (QD SPT) analysis.

### 2.2 Probing amyloid plaque accessibility by quantum dot tracking

TUSHI revealed that plaques come with a dense ring of cells, which might affect molecular access to the central core of the amyloid plaque. To study the accessibility of the amyloid plaques to macromolecules, we deployed photoluminescent quantum dots (QD) emitting at a wavelength of 655 nm. These particles are passivated by a PEG layer, and decorated with F(ab’)2 IgG (H+L, goat anti-rabbit) fragments to confer antibody-like diffusive behaviour [12]. Moreover, their diameter is comparable to biological macromolecules (*e.g.,* immunoglobulins, 10 – 20 nm) and nanocarriers (*e.g*., exosomes, 30 – 150 nm) present in the cerebrospinal fluid (CSF) [13]. We characterised the probes by imaging the dry size of the QD core and its organic shell (∼ 25 nm) using transmission electron microscopy of negatively stained nanoparticles (Supplementary Figure 3a). Additionally, we determined their hydrodynamic diameter (33.6 ± 1.2 nm) in physiological saline using dynamic light scattering (DLS) (Supplementary Figure 3b).

Incubating acute brain slices for 40 minutes, QDs penetrate the tissue and navigate the ECS (Supplementary Videos 6-7, respectively APP/PS1 and WT cortex). Their dimensions, isotropy, and photostability allow tracing their trajectories as they diffuse in the ECS. To locate amyloid plaques within the cortex in *ex vivo* carbogenated tissue (**Figure 3a**), we exploited the characteristic autofluorescence of plaque cores under confocal excitation with 405 nm light (Supplementary Figures 4 and 5) [14]. This allowed us to draw a concentric region of interest (ROI) around autofluorescent amyloid cores and to super-localise QDs (Supplementary Figure 6) inside the plaque, quantifying their accessibility and mobility in the regions of the amyloid core (*Amyloid*), cellular ring (*Ring*), and the immediate surrounding tissue (*Out*) (Figure 3b). While the cell ring was not visualised in the SPT experiments, we placed the ROI for it according to the TUSHI measurements, extending the radius of the amyloid core by a factor of 1.82 (Supplementary Figure 7).

**Figure 3.**
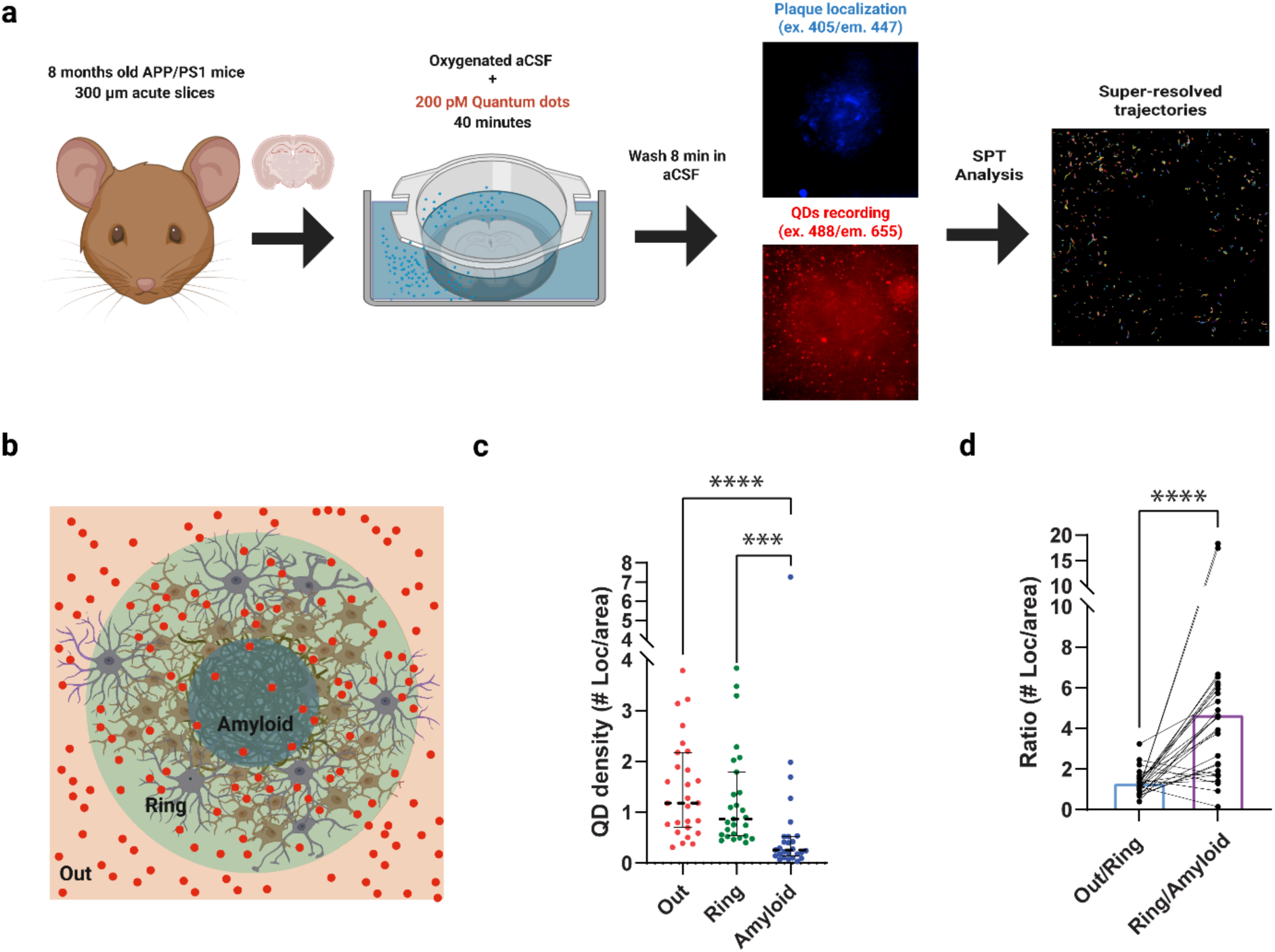
QD video-tracking enables the investigation of nanoscale diffusion dynamics within amyloid plaques. a - Schematic workflow of the QDs tracking in acute slices. 8-month-old females are sacrificed by cervical dislocation, and acute brain slices are then prepared in a bubbled ice-cold NMDG-based solution. After recovery, slices are incubated for 40min in oxygenated warm aCSF containing QDs (about 200 pM) and washed another 8 minutes in aCSF only. Slices are imaged in an upright microscope while perfused with aCSF at 35 °C under constant carbogen bubbling. Plaque was located using 405 excitation autofluorescence. QDs were imaged using 488 excitations and an emission filter. Analysis gauges the building of super-resolved trajectories of particles (see methods and Supplementary figure S6 for further information) b – Sketch of identified anatomical compartments in the proximity of senile plaques, consisting of 3 concentric ROIs: we determine QD (sparse red dots) behaviour as they move from the cortical parenchyma (Out), through cell dense envelope (Ring), until the centre of Aβ plaques (Amyloid). c - We found QDs localisations were scarcer inside the amyloid ROI in a plaque-dependent manner. To quantify this effect, we have quantified the particle density by ROI quantifying super-localizations per unit area (Out: Median=1.18 IQR=0.69-2.1, Ring: Median=0.88 IQR=0.54-1.79, Amyloid: Median=0.26 IQR=0.14-0.52, Friedman test Multiple comparisons: Out vs Ring p-value=0.17, Out vs amyloid p-value<0.0001, ring vs amyloid p-value=0.0002). NAPP/PS1 = 27 plaques, from 5 mice. d - We defined a penetrability ratio by dividing the amount of QD points in the outside ROI per area by the number of QD points inside the ring ROI per area. Higher ratio values mean that QDs cannot access the inner structure, while values close to 1 mean an even distribution of points. The Outside/Ring ratio gauges a median value of median = 1.14, IQR = 0.98−1.51, pointing that ring itself does not importantly impede the access to diffusing molecules, while Ring/Amyloid shows a 4-fold higher ratio of median = 3.97, IQR 1.75-5.96 (Wilcoxon test p-value < 0.0001). NAPP/PS1 = 27 plaques, from 5 mice. Created in BioRender. Bezard, E. (2024) BioRender.com/w96n099.

We measured QD densities in the three compartments for all recorded fields (Figure 3c). The density of QDs was highest in the surrounding tissue, intermediate in the cell ring and lowest inside the amyloid core (Figure 3c). To assess potential changes in particle behaviour between the different compartments, *i.e.,* to define whether a specific interface is preventing penetration, we defined two interfaces (*Out-Ring* and *Ring-Amyloid*) between pairs of adjacent ROIs. We calculated the QD density ratio for these pairs. Localisation density ratios between the surrounding cortical tissue and cell-rich ring distributed around 1 with low variability (Figure 3d), indicating that the interface between cortical neuropil and the cell ring does not appear to impose a barrier for molecular diffusion. Ratios of QD density within the ring *versus* the amyloid core were characterised by a 4-fold increase, indicating a low penetrability and/or retention by the amyloid core (Figure 3d).

This analysis indicates that (i) QDs penetrate the cell ring and the core of the amyloid plaques, but (ii) the overall plaque structure restricts macromolecule availability within the amyloid. We next moved to clarify how macromolecule availability might be affected by plaque diversity.

### 2.3 Different amyloid cores exhibit distinct penetrability

Extracellular amyloid in plaques exists in many shapes and sizes that could be related to multiple mechanisms of formation and toxicity [15]. Amyloid pathology is well characterised throughout AD progression, but the functional impact, temporal progression and neurotoxic effects of various types of Aβ deposits are still not fully understood [16]. Some authors have suggested that microglial processes in early filamentous plaques envelop individual fibrils, facilitating their compaction. Dense core plaques, in turn, can be considered as more mature and less toxic forms [10, 16–17]. Approaching amyloid plaque diversity from an extracellular perspective may offer insights into plaque-specific properties and their implications for disease progression and treatment.

We investigated whether differences in the density or morphology of amyloid plaques influence extracellular diffusion and core penetrability. Recent studies have introduced methods to quantify plaque phenotypes using Thioflavin-S (ThS) staining combined with circularity analysis. This approach yields quantitative and qualitative indicators of plaque size diversity, offering robust and consistent results in both human tissue and genetic mouse models (see Methods for details) [10, 18]. We applied the same analysis to the APP/PS1 plaques (**Figure 4a**), assigning a circularity value to each plaque. Since ThS binding may interfere with QD penetration and signal, we developed a post-hoc labelling protocol: amyloid cores were first localised (based on their autofluorescence under UV light excitation), their coordinates recorded, and QDs tracked. After several recordings, the slice was incubated with 0.001% ThS in aCSF within the recording chamber. Finally, z-stacks of the previously studied (QD tracking) plaques were captured.

**Figure 4.**
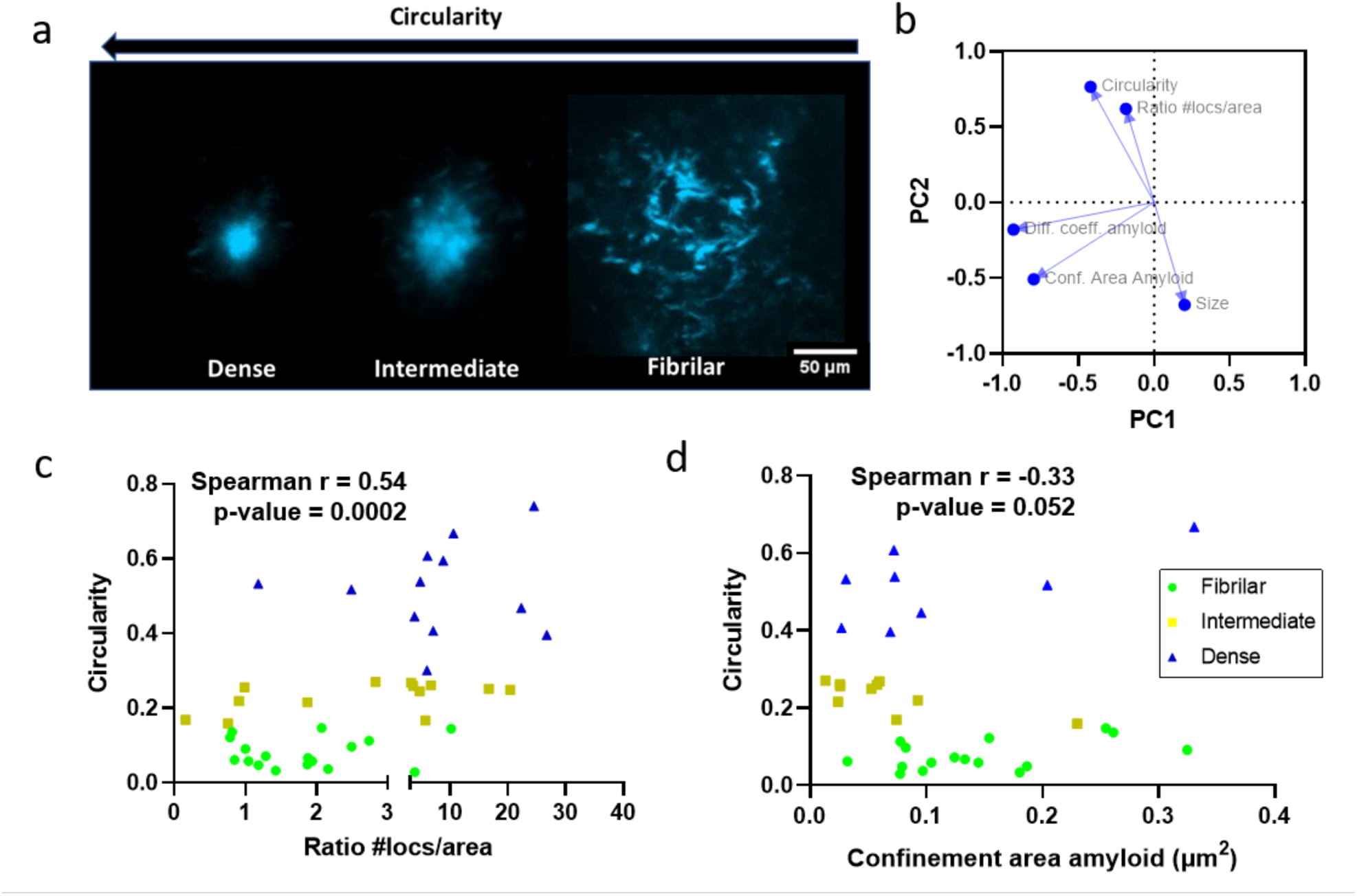
QD penetration depends on plaque core phenotype. a – After QDs recording, each plaque ThS signal was associated with a circularity value as a proxy of plaque phenotype (see methods) b - Principal Component Analysis (PCA) loadings plot shows association between circularity and QD density ring/amyloid ratio while suggesting a lack of linear correlation with rheological parameters (PC1 total variance = 34.9%, eigenvalue = 1.746: PC2 total variance = 34.4%, eigenvalue = 1.720). c - Scatter plot of Circularity versus ratio #locs/area (amyloid impenetrability) shows that as plaque densifies, penetration becomes more difficult. The relationship correlates significantly (Spearman r =0.54, p = 0.0002). Amyloids are categorised as previously defined [10] (fibrillar = 0.00–0.14, intermediate 0.15–0.28 compact > 0.28). d - We observed a strong trend for negative correlation between circularity and confinement area, suggesting that as plaques become denser, movement within them is progressively restricted (Spearman r =-0.33, p = 0.052). n=41 plaques from 4 mice.

To investigate the relationships among the experimental variables, we performed Principal Component Analysis (PCA), which revealed a clear association between circularity and impenetrability of the amyloid (QD ratio #locs/area, as defined in Methods). However, all diffusional parameters were oriented perpendicularly to circularity, suggesting a lack of linear correlation (Figure 4b). We correlated and plotted circularity and QD ratio #locs/area (Figure 4c). Circularity significantly correlates with QD ratio, showing that more circular (denser) plaques exhibit reduced QD penetration, consistent with limited antibody accessibility reported by others [10]. Despite the lack of a strong linear relationship in the PCA, we found an almost significant negative correlation (Figure 4d) between circularity and confinement area (see Methods and **Figure 5a** for definition), indicating that as plaques densify, both penetration and movement within them become increasingly restricted.

**Figure 5.**
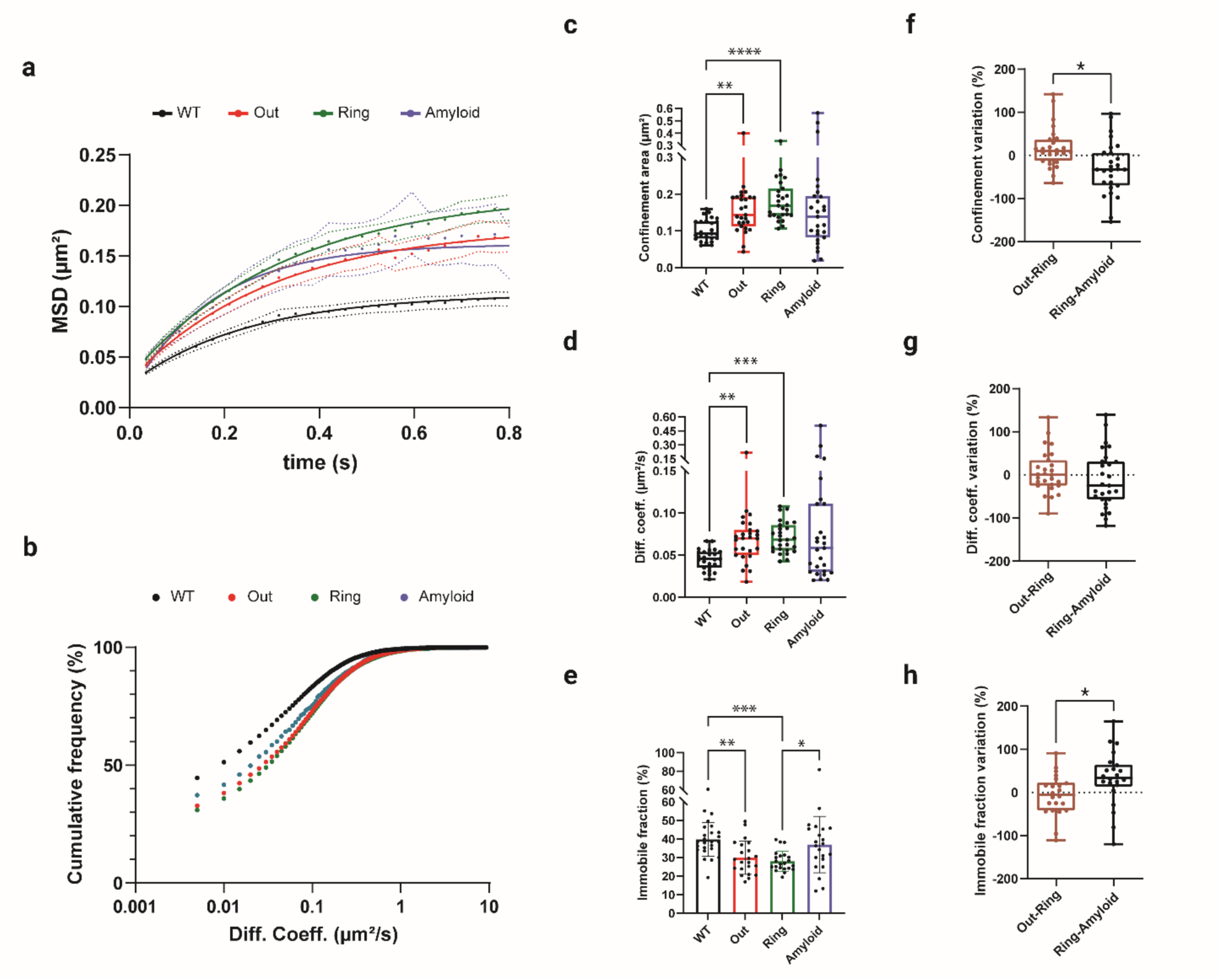
The rheological properties of the ECS differ between wild-type tissue, around and within plaques. a - The mean square displacement (MSD) curve suggests limited particle diffusion within the ECS. The confined area was estimated to indicate particle exploration (Fitted curve plateau: WT = 0.1123, Out = 0.1792, Ring = 0.2127, Amyloid=0.1617). b. The cumulative frequency distribution of all particle velocities (log scale) was left-shifted (slower mobility) in the WT (black) compared to the three compartments (coloured curves) around and within plaques. c-Comparison of confinement as a proxy of explored ECS area at MSD curve plateau (0.6 s – 0.8 s). In the surroundings of amyloid deposits, but not within it, a significant increase in confinement was observed compared to WT cortices (WT: Median = 0.093, IQR = 0.079-0.125, Out: Median = 0.143, IQR = 0.113-0.192, Ring: Median = 0.169, IQR = 0.143-0.215, Amyloid: Median = 0.139, IQR = 0.082-0.194; Kruskal-Wallis test (Dunn’s multiple comparisons correction) p-values: WT vs out = 0.0054, WT vs ring<0.0001, WT vs amyloid = 0.069, Out vs Ring = 0.58, Out vs amyloid >0.99, Ring vs amyloid = 0.08). d - Likewise, the medians of diffusion coefficient indicate higher particle velocities in the AD model brain than in WT (WT: Median = 0.045, IQR = 0.035-0.053, Out: Median = 0.07, IQR = 0.050-0.08, Ring: Median = 0.068, IQR = 0.055-0.085, Amyloid: Median = 0.058, IQR = 0.03-0.11, Kruskal-Wallis test (Dunn’s multiple comparisons tests, p-values): WT vs out = 0.0021, WT vs Ring = 0.0002, WT vs amyloid 0.075, Out vs Ring >0.99, Out vs amyloid >0.99, Ring vs amyloid = 0.54). NWT = 25 FOV, from 4 mice; NAPP/PS1 = 27 plaques, from 5 mice. e - Quantification of the immobile fraction of particles highlights an overall smaller fraction in the cortical neuropil (Out) of AD mice compared to controls, with an even greater difference in the cell-dense ring (WT: Median = 38.4, IQR = 34.5-45, Out: Median = 28.2, IQR = 23.1-37.2, Ring: Median = 26.7, IQR = 24.2-30.9, Amyloid: Median = 36.9, IQR = 28.1-45.6; Kruskal-Wallis test (Dunn’s multiple comparisons) p-values: WT vs out = 0.0043, WT vs ring = 0.0003, WT vs amyloid >0.99, Out vs Ring >0.99, Out vs amyloid >0.23, Ring vs amyloid = 0.036). NWT = 25 FOV, from 4 mice; NAPP/PS1 = 22 plaques, from 5 mice. f - Paired analysis demonstrates that relative differences in explored area values during the transition from out to ring distributed around zero, in contrast with the transition from cell ring to amyloid plaques, where they assumed negative value (Out-Ring: Median = 10, IQR =-13-37, Ring-Amyloid: Median = -33, IQR = -70-6; Wilcoxon pairs signed rank test, p-values = 0.01). g – Paired analysis of diffusion coefficient variation in the transition between the two adjacent areas showed a comparable trend, indicating that median diffusion also decreases from ring to amyloid. Such decrease is not statistically significant from that between out and ring (Out-Ring: Median = 0, IQR =-25-34, Ring-Amyloid: Median = -25, IQR = -57-31; Paired t-test, p-values = 0.27). NAPP/PS1 = 27 plaques, from 5 mice. h – Finally, the extent of trapping within the plaques was evaluated through relative variation of the immobile fraction in successive compartments. Again, the passage from ring to amyloid resulted in a greater variation (increase in this case) in the number of immobile particles (Out-Ring: Median = -5.5, IQR =-42-23.5, Ring-Amyloid: Median = 34, IQR = 13.7-64.7; Paired t-test, p-values = 0.044). NAPP/PS1 = 27 plaques, from 5 mice.

Finally, a strong negative correlation exists between highly circular plaques and area, supporting the hypothesis proposed by Condello and colleagues that denser amyloid plaques represent an advanced and more compact stage of fibrillar plaques [17]. However, plaque size alone is not a reliable predictor of the penetrability of the amyloid core (Supplementary 8).

### 2.4 Rheological ECS properties within and around amyloid plaques

Beyond penetrability, extracellular rheological parameters are of great relevance when studying the ECS. Mean square displacement (MSD) analysis provides rheological information, i.e. diffusionality and dimensions, and ECS nanoscale properties from SPT experiments in biological media [19]. We plotted MSD curves and diffusion coefficient distributions of super-resolved QDs trajectories within and around amyloid plaques over time. On the one hand, diverse curve plateaux were observable at a lag time of 0.4 s, indicating that particle displacement within the ECS was limited to a typical explored area (Figure 5a). On the other hand, the distribution of diffusion coefficients in the ECS exhibited marked differences between experimental conditions (Figure 5b). To quantify these differences statistically, we extracted (from MSD plateaux between 0.6 and 0.8 s) the average confinement area to estimate the space the diffusing particles explored within a given time interval. From both MSD curves and the distribution of confinement values, a dramatic increase in the confinement area was observed in the cortex of APP/PS1 mice compared to age-matched wild-type (WT) controls (Figure 5c). By contrast, the distribution of confinement values within the amyloid core was not significantly different from WT tissue. Still, the distribution was much wider, likely reflecting other physico-chemical differences and amyloid phenotype influence, with a prevalence of extreme values, indicating a significant change in particle mobility within fibrillary aggregates (Figure 5c). QD dynamics were further addressed by the analysis of diffusion coefficient distribution (see Methods). We set a threshold value (0.005 µm²/s) to study separately the mobile and immobile fractions of particles within the ECS at each level [20]. The median diffusion coefficient of the mobile fraction was higher outside the amyloid plaque and in the cell ring in the AD model than in age-matched WT cortical tissue (Figure 5d). Although not statistically significant, the distribution of the diffusion coefficient was wider for the amyloid region, suggesting alterations in the ECS. Interestingly, both the cortical neuropil (*Out*) and cell ring (*Ring*) of APP/PS1 mice exhibited a significantly smaller fraction of immobile particles compared to WT tissue (Figure 5e). Among the few particles that had penetrated the amyloid core, a sizable percentage was immobile, more so than in the surrounding ring (Figure 5e).

We deepened our confinement analysis by pairing values for the three regions (*Out*, *Ring* and *Amyloid*) obtained from the same field of view. Again, we focused on the two interfaces (*Out-Ring* and *Ring-Amyloid*) for which we calculated distributions of their differences expressed as relative change (%) with respect to the average value of each of the two adjacent regions (see Methods) for all parameters as above. Interestingly, the explored area, diffusion velocities, and immobile fraction displayed substantial variations between the two interfaces. For these three parameters, the percentage of relative change from the outer zone to the ring (Out-Ring) is distributed around zero, which means no variation among neighbouring regions. By contrast, at the ring-to-amyloid interface (*Ring-Amyloid*), we observed marked differences between the compartments. Confinement area and diffusion coefficient variations settled around a negative value (i.e., rheological parameters became lower inside the plaques), while the immobile fraction variations were elevated inside the plaques (Figure 5f-g-h).

Altogether, our approach unveiled substantial nanoscale alterations in the rheological properties of APP/PS1 mouse cortex compared to age-matched WT. These pathological changes resulted in the unexpected diffusion behaviour of particles proximal to Aꞵ plaques, where mobility was facilitated even within the cell-dense ring. In parallel, we found a reduction in the availability of nanoparticles within the plaque. Finally, this reduction was more accentuated in the transition from ring to amyloid.

Finally, there is a current controversy regarding the *in vivo* existence/extent of active flow in the brain parenchyma [21]. To test if our ex vivo model resembles in vivo diffusion, at least regarding MSD shape, we performed *in vivo* tracking of QDs using 2-photon microscopy in WT mice. After the topical application of QD-containing solution on the mouse cortex, followed by rinsing, we recorded *in vivo* trajectories in depths ranging from 20-40 μm. The analysis shows that most particle diffusion exhibits a restricted/sub-diffusive profile that resembles our observations in *ex vivo* settings with a clear MSD plateau after 1.2 seconds, supporting the validity of ex vivo experimentation (Supplementary Figure 9 and Supplementary Videos 8-9).

### 2.5 Cortical Amyloid plaques exhibit a degraded extracellular matrix

Despite the highly crowded environment in the plaque ring, ECS rheological parameters (Figure 5c-d) are consistent with a facilitated diffusion microenvironment. Such a counterintuitive observation led us to question the integrity of the ECM, a major diffusional regulator of the ECS [22]. ECM alterations have emerged as a common phenomenon in several neurological conditions [23] and amyloid animal models [24], being directly related to the observed increase in ECS diffusion [5b]. We first studied immunolabelled glial cells (IBA-1 and GFAP+S100) and hyaluronic acid (labelled by Hyaluronic acid binding protein: HABP) around the autofluorescent amyloid core. While the ring is primarily composed of astrocytes and microglial cells [25], we report that amyloid plaques present a highly degraded ECM, with a decreasing HABP signal from the outer ring to the core of the plaque (**Figure 6a**). For consistency, we used the ROI system defined in previous figures (Figure 6b). ECM changes were quantified by assessing the surface occupied by HABP (Figure 6c). To rule out a decrease in total signal due to a lower ECS volume fraction, we also quantify the fractal dimension [26] of the HABP thresholded signal, to estimate matrix interconnectivity and organisation (Figure 6d), confirming a significant decrease. The dramatic decrease in hyaluronic acid content in the ring and amyloid supports the observation of an increased QD local diffusion (with the immobile fraction decreasing in the ring) despite the apparent crowded cellular environment. Conceivably, it provides an explanation for why extreme diffusional values are found in the amyloid, i.e., QDs penetrating the amyloid meshwork of proteins/organelles would either stick (immobile) or diffuse (highly mobile) in an amyloid devoid of ECM.

**Figure 6.**
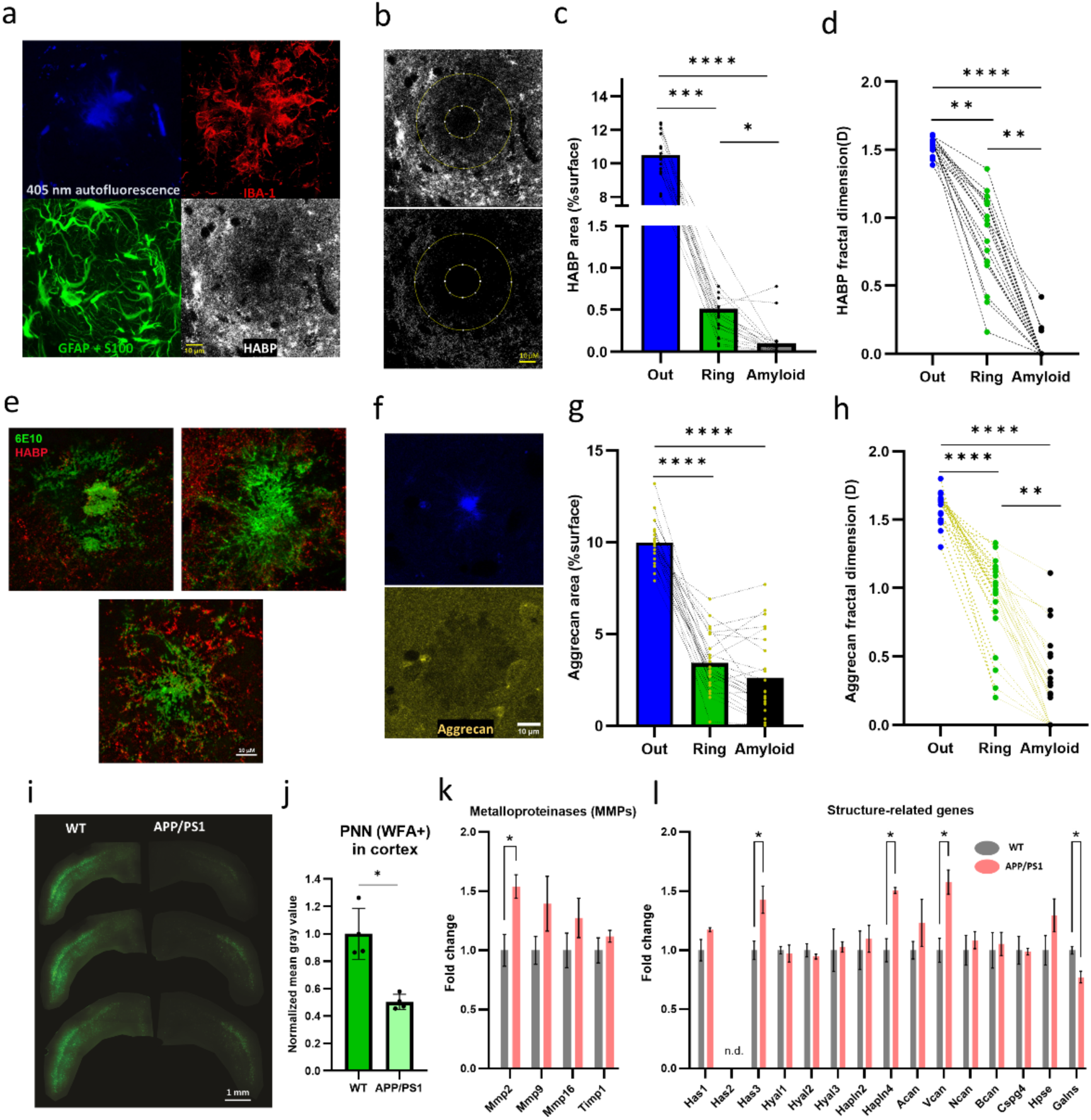
The APP/PS1 mouse presents a highly modified extracellular matrix. a - Amyloid plaques are surrounded by activated astrocytes (GFAP) and microglia (IBA1). Importantly, they also present a highly disrupted Hyaluronic Acid (labelled by HABP) matrix scaffold. b - Matrix thresholding and skeletonisation. Matrix is practically non-existent in the amyloid ROI and is severely degraded in the ring ROI. For segmentation, we applied the same ROIs as in Figure 3. c - Quantification by percentage of HA signal per unit area (%) Ext. Median = 10.50, IQR = 9.52-11.90; Ring. Median = 0.371, IQR = 0.253-0.566; Amy. Median = 0.0175, IQR = 0.0027-0.0750, Kruskal-Wallis test p < 0.0001, (Dunn’s multiple comparisons correction) p-values: Out vs Ring = 0.0004, Out vs amyloid <0.0001, Ring vs amyloid = 0.021. d - Fractal analysis (Ext. Median = 1.56, IQR = 1.52-1.58; Ring. Median = 0.96, IQR = 0.66-1.11; Amy. Median = 0, IQR = 0, Kruskal-Wallis test p < 0.0001 (Dunn’s multiple comparisons correction) p-values: Out vs Ring = 0.0013, Out vs amyloid <0.0001, Ring vs amyloid = 0.0022. N =18 plaques, from 4 mice. e - Expansion microscopy of amyloid β-immunolabelled plaques (6E10 antibody) and hyaluronic acid (HABP) permits the visualisation of the voids and ECM state inside the plaques. f – Aggrecan was labelled and analysed the same way as HA g - Quantification by percentage of aggrecan signal per unit area (%) Ext. Median = 9.90, IQR = 9.40-10.60; Ring. Median = 3.10, IQR = 2.36-4.8; Amy. Median = 1.73, IQR = 1.21-4.55; Kruskal-Wallis test p < 0.0001 (Dunn’s multiple comparisons correction) p-values: Out vs Ring = <0.0001, Out vs amyloid <0.0001, Ring vs amyloid = 0.57. h - Fractal analysis (Ext. Median = 1.64, IQR = 1.54-1.66; Ring. Median = 0.99, IQR = 0.85-1.12; Amy. Median = 0.215, IQR = 0-0.473, Kruskal-Wallis test p < 0.0001 (Dunn’s multiple comparisons correction) p-values: Out vs Ring = < 0.0001, Out vs amyloid <0.0001, Ring vs amyloid = 0.031. i - PNN were labelled with WFA and quantified in the sensorimotor cortex. j - There is a significant decrease in WFA intensity in APP/PS1 mice (Mann-Whitney test p-value = 0.029, n = 4 per group). Each data point represents the average grey value per mouse, calculated from three anterior-posterior cortical slices corresponding to the sensory-motor region. k-l mRNA expression analysis of ECM-related genes in cortex homogenates of WT and APP/PS1 mice. APP/PS1 mice present an altered expression pattern of several key regulator components of ECM. j - Metalloproteinases (MMPs) expression levels show an increase in the APP/PS1 mouse (MMP2 p-value = 0.032). k - Matrix structural-related genes present altered expression patterns, with upregulated (Has3 p-value = 0.033, Hapln4 p-value = 0.015, Vcan p-value = 0.019) and downregulated expression (Galns p-value = 0.022). Error bars represent mean ± SD. Two-tailed Student’s t-test, n = 3 mice. The source data is provided as supplementary.

We next used expansion microscopy (Figure 6e) of amyloid β-immunolabelled plaques to visualise the voids and ECM state (labelled again by HABP) inside the plaques, a complementary approach to the earlier work on the tissue-level impact of amyloid fibres in the 5xFAD mouse model of AD [27]. Amyloid appeared susceptible to an expansion, resulting from the insertion of the sodium acrylate between amyloid fibres, i.e., in the ECS, likely filled by the dye in the TUSHI and nanoparticles in the QD experiments, respectively.

In agreement with the classic description of diffuse and dense plaques discussed before [28], we observed that protein aggregates present a high degree of structural heterogeneity in ECS and ECM between the fibres (Figure 6e), consistent with the variability in the nanoscopic single-particle tracking data. To expand our ECM analysis, we imaged the proteoglycan aggrecan, one of the main components of the ECM [23]. Following the same protocol and analysis for HABP, we also found a decrease in aggrecan signal, albeit more modest, pointing out that matrix disruptions also affect the proteoglycan component of the matrix (Figure 6g-h).

Finally, we aimed to assess if matrix disruption can be found beyond plaques. Crapser et al. (2020) reported that PNNs are extensively lost in AD patients and the 5xFAD mouse model in proportion to plaque burden [29]. Following a similar approach, we labelled PNN with WFA and observed a significant decrease in WFA-binding PNN in the APP/PS1 sensory-motor cortex (Figure 6i-j) pointing towards a structural disruption or changes in the “sulfation code” in these key structures [30]. To further understand matrix alterations in APP/PS1 mice beyond immunostaining, we examined mRNA expression of genes associated with matrix degradation, structure, and metabolism in cortex homogenates of WT and APP/PS1 mice. We studied the expression of the matrix metalloproteinase family (MMPs) (Figure 6k), given its crucial role in matrix regulation and degradation in health and disease [23, 31]. Several groups have reported increased endopeptidase levels in amyloid mouse models [32] and human AD [31], suggesting their involvement in clearing Aβ. Relevant to this work, MMPs degrade and regulate the turnover and remodelling of ECM proteins [33], making them prime candidates for causing disease [23]. We found a significant increase in MMP-2 mRNA levels and a general increase in the other mRNAs coding for proteins of the family. The rise in Tissue Inhibitor of Metalloproteinases 1 (TIMP-1) might reflect a counter-balancing attempt to dampen the consequences of MMP upregulation. It suggests that the over-expression of MMPs could contribute to the reported degradation of the ECM, conceivably through microglial activation [34]. Next, we focused on the expression of matrix structural components (Figure 6l). Hyaluronan synthase 3, Hyaluronan/Proteoglycan Link Protein 4, and Versican showed increased mRNA levels in APP/PS1 mice and Chondroitinase mRNA level is downregulated, probably in a compensatory attempt to regulate or repair digested components. In conclusion, we show that amyloid pathology is also reflected in a highly altered expression pattern of several key matrix components/regulators.

### 2.6 Rheological ECS properties in the amyloid pathology-bearing cortex

The collected results provide a comprehensive picture of molecular diffusion in the ECS in and around amyloid plaques. Remarkably, we found that amyloid brain rheology around plaques exhibits significant differences compared to WT (Figure 5). Besides, we make the point that ECM alterations are widespread in amyloidosis. This prompted us to investigate further whether an amyloid brain undergoes global alterations in ECS rheological properties, especially an increased diffusion, using a reporter agnostic of the proximity of amyloid plaques.

Single-walled carbon nanotubes (SWCNTs) represent a complementary SPT technique using slowly-moving near-infrared reporters, *i.e*., working in the “transparent” spectral window of biological tissue [35]. Besides local diffusion coefficients, our analysis based on local restricted theory [36] can provide access to the spatial dimensions of the ECS, similar to cryofixed EM [5b] and super-resolution shadow imaging (SUSHI) [37]. We recently used SWCNTs as super-resolution imaging probes to report on the local nanoscale organisation and diffusivity of the ECS in young [36] and adult [35] living brain tissue. We also demonstrated that several mouse models of Parkinson’s disease were characterised by increased nanoscale diffusion [5b, 38]. Applying this SPT technique to our APP/PS1 mice, we now unravel a heterogeneous cortical ECS, where local diffusion properties change over distances of just a few micrometres and are effectively shaped by brain ECS geometry and composition (Figure 7a). Direct comparison between age-matched WT and APP/PS1 mice showed an overall increase in local diffusion [35] (D_inst_, Figure 7b). Concomitant with the alterations in local diffusion, we also report a widening of the ECS, i.e., an increase in local channel width based on the analysis of local nanotube trajectories [35] (Figure 7c). In agreement with restricted diffusion theory [39], local diffusion and channel width present a weak positive correlation (Figure 7d), suggesting particle diffusion scales with ECS width. However, the modest correlation coefficients indicate that diffusion is a multifactorial parameter, which depends not only on size and geometry. The correlation increases in the amyloid model, pointing to other factors than wider ECS spaces behind the observed increase in diffusion. It raises the question of how a dysregulated ECM affects diffusion in different brain regions. The nuanced relationship between diffusion and channel width underscores the complexity of their interplay. Finally, the SWCNTs and QDs experiments provide converging evidence that amyloid pathology leads to a more diffusive ECS in the cortical neuropil of APP/PS1 mice compared to WT controls (Supplementary Figure 10).

**Figure 7.**
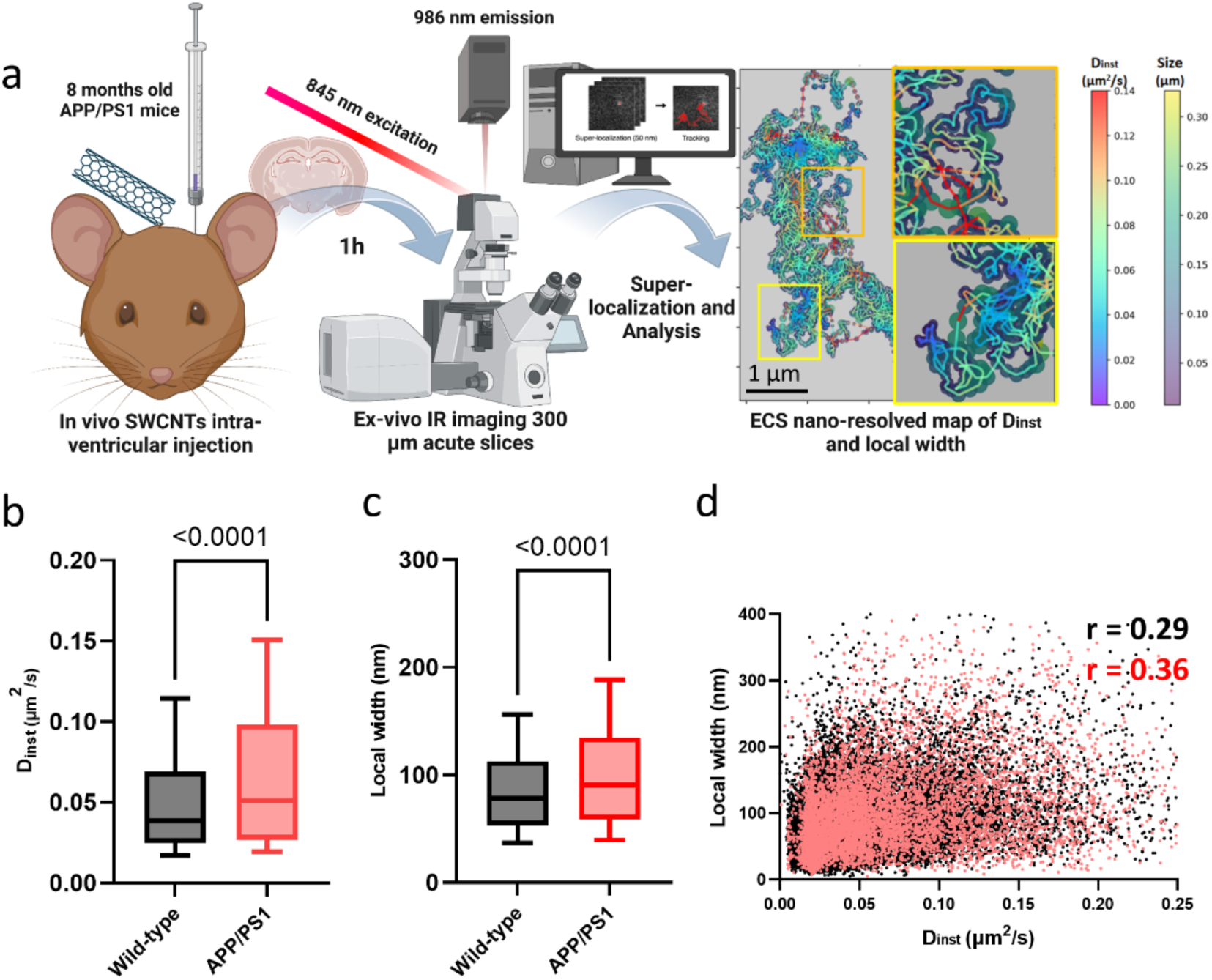
SWCNT tracking reveals a diverse nanoscale ECS and increased local diffusion and ECS width in amyloid pathology. a - SWCNTs are in vivo intra-ventricularly injected in 8-month-old mice and allowed to diffuse into the brain for 1 h. Acute brain slices were prepared in ice-cold NMDG-based solution and imaged in aCSF at 37 °C under carbogen bubbling. A laser emitting at 845 nm was used to excite (6,5) SWCNTs at their phonon sideband, and their emission peaked at 986 nm. They were filtered using a long pass filter before being imaged and recorded with a water-cooled SWIR InGaAs camera. Individual SWCNTs were super-localized with a 2D Gaussian fit, and coordinates were linked to reconstruct individual trajectories. Each SWCNTs trajectory provides a nanometric-resolved map of local instantaneous diffusion coefficients (µm^2^/s) and local widths (nm) (see supplementary Figure 12 for further details). ECS Dinst and local size maps reveal a heterogeneous ECS in the brain, with properties that change significantly within a few nanometers. Scale bar = 1 µm. b - Distributions of local Dinst values increase in diffusion values in the APP/PS1 model (Dinst (µm^2^/s) WT Median = 0.038, IQR = 0.025-0.069, APPPS1 Median = 0.051, IQR = 0.027-0.98, Kolmogorov-Smirnov test p-value < 0.0001). c - Distributions of local ECS widths (nm). We show an increase in local width values in the APP/PS1 model (local width (nm) WT Median = 78, IQR =53-113, APPPS1 Median = 90, IQR = 59- 135, Kolmogorov-Smirnov test p-value < 0.0001). d - There is a positive but weak spatial correlation between size and Dinst, increasing in pathology (p < 0.0001 for both groups, Pearson’s r = 0.29 WT, 0.37 APP/PS1). NWT = 4 mice; NAPP/PS1 = 4 mice. Created in BioRender. Bezard, E. (2023) BioRender.com/i55x918.

## 3. Discussion

The study employs three advanced imaging techniques, including TUSHI and QD / SWCNT single-particle tracking, to investigate the morphology and diffusionability, i.e. the rheological properties, of the ECS in and around cortical amyloid plaques, formed by an amyloid core surrounded by a dense ring of cells in a transgenic mouse model of AD. The QD-SPTinvestigation revealed an increased permeability of the ECS within and around these plaques, but low amyloid core penetrability, demonstrating how cortical ECS is remodelled by amyloid pathology and how amyloid plaques display exploitable features for future therapeutic development. These rheological alterations are likely key to understanding the disease. They may contribute to the spreading of the pathology and impact the delivery of therapeutics, as both phenomena are highly dependent on the anatomical fine structure of the tissue and its physico-chemical properties [39].

Despite its pivotal role in essential processes like synaptic and extra-synaptic communication, extracellular ionic homeostasis, drug delivery, and metabolic waste clearance [22], the properties of the ECS have been predominantly overlooked in the scientific literature, partly because of the lack of suitable techniques to address this challenge. Few methods have been developed to measure extracellular diffusion, including diffusion-weighted magnetic resonance imaging (DWI-MRI), electrochemical methods, and optical imaging [4, 40]. New optical imaging approaches have since been developed (reviewed in [41]). Among them, super-resolution shadow imaging (SUSHI) was designed to address the microanatomical structure of the ECS in living brain tissue with nanoscale spatial resolution [22a]. Our TUSHI methodology, which is a diffraction-limited variant of it, enabled visualisation of amyloid plaques deep inside brain tissue. This achievement holds significant importance in the field as it offers a non-invasive means to explore and comprehend the distribution and dynamics of amyloid plaques in their native environment, facilitating the analysis of the intricate interplay of amyloid plaques with the surrounding tissue. Using TUSHI, we observed that small dyes can penetrate the very centre of the amyloid plaque. While we could not resolve the internal plaque structure with TUSHI nor obtain ECS volume fractions, the images show that the amyloid plaques are not an impenetrable agglomeration of proteins and organelles but rather a porous-like mesh formed by amyloid fibrils, sugars, dystrophic neurites and cellular components that can be penetrated and filled by small molecules, hence the apparent “void” in the diffraction-limited images (Figure 1, Supplementary videos 1-5). In addition, it strikingly highlighted the known large and dense ring of cells that surround the amyloid core like a mantle, increasing by a factor of 25 the effective volume occupied by the plaques. Notably, it also allowed us to define the ROI dimensions required for further exploration with single-particle tracking.

Proteomic studies that report up to thousands of proteins illustrate the complex composition and crowdedness of amyloid plaques [42]. Still, although protein, lipidic and organelle content have received much attention [9], our understanding of how small and larger molecules diffuse in and around amyloid plaques is rudimentary despite its relevance for therapeutics, notably the amyloid-targeting immunotherapies [43]. It is striking that the most advanced quantitative systems pharmacology (QSP) models that describe the non-linear progression of amyloid ß pathology and the pharmacological actions of the amyloid ß-targeting antibodies do not consider the penetrability of amyloid plaques or their capacity to accumulate exogenous substances [44]. While the movement of small molecules and ions is relatively easy to model, understanding how larger objects diffuse within the ECS and interact with ECM components is more challenging due to the multifactorial dependency of extracellular diffusion [22a]. The single particle tracking methods utilise fluorescent probes diffusing in the ECS, allowing for real-time monitoring of their trajectories and speeds and for accessing the penetrability of plaques and overall ECS rheology.

QDs and SWCNTs provide complementary information owing to their specific optical properties and physical dimensions. On the one hand, spherical QDs penetrate the acute brain slice upon incubation. ECS rheology (diffusionality and dimensions) can thus be measured in live tissue with high throughput (many moving particles per field of view). On the other hand, SWCNTs allow thorough exploration of local geometry based on their elongated shape (much thinner but much longer than QDs) and their much better photostability than QDs. More precisely, we took advantage of the fact that the transverse local ECS dimensions laterally constrain the diffusion of the SWCNTs and, therefore, that these dimensions can be locally measured by analysing the local excursion of the SWCNT [36]. In other words, SWCNTs can give access to both local ECS dimensions and local diffusivity. However, SWCNTs do not penetrate the tissue as well as QDs do. The *in vivo* ICV approach, although elegant, provides a much lower throughput than QDs. This means that, practically speaking, it is complicated to find SWCNTs inside a plaque and even harder to find several in the same field of view to compare the values. In conclusion, the QD method provides shorter trajectories but much higher throughput, while SWCNTs provide better photostability and give access to local dimensions. Besides, we believe that replicating the finding of an increased diffusion in APP/PS1 mice with different probes (which have distinct dimensions and surface chemistry) greatly strengthens our results (Supplementary Figure 10).

When examining diffusion in the brain, the nature and extent of different types of material and fluid flow become an important question. The emergence of the glymphatic system has introduced the idea of active flow in the brain [45]. This type of flow would not be Brownian or passive. Instead, according to the glympathic system hypothesis, there is an active process that directs and powers fluid flow along specific pathways. In the absence of blood flow and intracranial pressure, the *ex vivo* conditions do not replicate fully the diffusion/flow situation found *in vivo*. There is, however, an ongoing controversy regarding the extent of active flow, especially in the parenchyma, where our experiments take place. Many researchers believe that the hydraulic resistance in the narrow channels of the ECS is too high for any bulk flow to occur, even with significant pressure [21]. To address this in our model, we performed *in vivo* experiments using a 2-photon microscope with enough temporal resolution to track and reconstruct QD trajectories. Our results provide crucial proof-of-concept for the existence of restricted diffusion *in vivo*, which seems to be predominant in the parenchyma, supporting the robustness of our *ex vivo* model. However, we must concede that our *in vivo* recordings are far from optimal for super-resolution SPT analysis. A relatively big pixel size and lower frequency restrict the accuracy of measurements. Besides, we are limited to the first layer of the cortex, which is likely not representative of the wider cortex. This means that despite the important proof of concept these measurements represent, for the moment, we still rely on *ex vivo* measurements for quantification.

Besides, our *ex vivo* approach offers an additional advantage. By disconnecting the presence of putative flows from ECS remodelling, we characterise constitute intrinsic properties of the system. The nanoscale SPT approach measurements provide a “mapping” of the local ECS (local Dinst can be considered analogous to local viscosity), which is relevant irrespective of any flow considerations.

Regarding probe selection, functionalised QDs represent a powerful approach to mimic antibodies in size and cell interactions. Although amyloid plaques exhibit a dense ring of cells, QD diffusion in this ring is not impaired. It even appears slightly facilitated, possibly reflecting a remodelled ECS after a loss of ECM. However, the influence of ECM on local hydraulic properties remains correlative and not causal in our experiments.

An interesting point concerns QD density: despite matrix disruption in the “ring” region of the plaque, we report a similar QD density outside the plaque. Even if rheology is facilitated by a dysregulated matrix, cellular components reduce the available extracellular space and, thus, the likelihood of finding a particle and QD accumulation. This equilibrium between low resistance and low available space likely renders the final QD density in each region. In the amyloid core, diffusion and access are restricted in a plaque-dependent manner, likely due to size-selective barriers: We show amyloid core penetrability is highly dependent on plaque compaction, defined by circularity [10], rather than plaque size, rendering certain plaque phenotypes significantly more accessible to antibodies than others. Several studies have shown microglia mutations and relative populations affect the amount the relative number of each kind of plaque, affecting toxicity[10, 46]. This suggests that a temporal window for successful immunotherapy against plaques exists before microglial-mediated plaque compaction might render treatments less effective. After such a stage, immunotherapy should be assisted by microglial-directed therapy (cf. Lixisenatide reports on the suppression of microglial reactivity) [47].

Beyond plaque accessibility, our data suggest a cortex-wide reorganisation of the ECS with widening spaces and increased diffusion. The widening of brain ECS in AD has been implicated in the dissemination and propagation of misfolded/aggregated Aβ and tau proteins, allowing the misfolded proteins themselves, or extracellular vesicles containing them, to propagate from regions of initial pathology to other brain areas later affected during disease progression [48]. It can disrupt physiological synaptic transmission and neuronal communication [49]. This disruption, in turn, may contribute to the spread of pathological proteins by altering the microenvironment of neurons and facilitating their vulnerability to protein aggregation [50]. The ECS also plays a crucial role in interstitial fluid dynamics, which is essential for the clearance of metabolic waste products, including Aβ and tau. ECS width changes can impact interstitial fluid flow, potentially leading to impaired clearance and the accumulation of pathological proteins, as suggested by others [51]. Finally, ECS widening is often associated with neuroinflammatory responses in AD [52], which, in turn, may exacerbate the spread of pathological proteins by promoting cellular damage and dysfunction, creating an environment conducive to the propagation of Aβ and tau pathologies. These hypotheses can now be thoroughly tested using the developed technologies.

Further exploring how plaque composition, maturity and morphology influence drug penetration could inform personalised treatment strategies tailored to the stage of AD progression. Here, however, we used a transgenic mouse model of AD, the APP/PS1 mouse model. Since 1995, more than 100 transgenic mouse models of AD have been developed expressing high levels of mutant human amyloid precursor protein showing varying degrees of symptoms, pathophysiological processes and plaque formation [53]. While the Tg2576, APP23, APP/PS1 and 5xFAD lines are popular in the literature, their validity remains debated [54]. The pathological changes in APP/PS1 mice show several similarities with AD: the abundant age-dependent severe neuropathology [55] is associated with global brain atrophy [56], decreased glucose metabolism in the hippocampus [57] and complex AD-like cognitive [58] as well as non-cognitive behavioural manifestations [59]. The model inevitably also presents discrepancies with AD. For example, it is based on overexpression of APP, which does not happen in human AD (a limitation that has led to the third generation of mouse AD models, the KI lines [60]), while hyperphosphorylated tau remains at low levels, is distributed in a different way than in human AD patients and does not form neurofilaments [61]. The lack of tau pathology is held responsible for the limited neuronal loss and lower amyloidogenic processing of the amyloid precursor protein (APP) as compared to humans [62].

This study underscores the importance of the dynamic nature of AD pathology for developing improved therapeutic strategies. Not only does it offer a screening platform for measuring the penetrability of said experimental therapeutics in amyloid plaques, but it also paves the way for elaborating nanotechnology-based approaches, such as targeted drug delivery systems or modulators of ECS dynamics, to improve drug penetration and efficacy for advancing AD therapeutics. However, a limitation of the current setup is the simultaneous shadow imaging and SPT data acquisition, preventing the parallel study of the same plaque. Technically, integrating shadow imaging, particularly in combination with STED, alongside SPT, represents a challenge. This approach would allow for concurrent analysis of cellular architecture and dynamic processes, offering deeper insights into the relationship between mechanical and structural features and their impact on biological functions. Specifically, it would be highly valuable to link plaque rheology and penetrability with amyloid aggregation and different amyloid morphologies.

The QDs-SPT technique we propose could answer other biologically meaningful questions. For instance, tumours represent compelling structures for extracellular diffusion measurements using our QDs SPT protocol. Like amyloid plaques, tumours present a particular tumour microenvironment (TME) characterised by altered ECM and diffusion [63]. ECM remodelling—marked by increased hyaluronic acid, tenascin-C, tissue stiffness, and MMP activity—supports tumour cell migration, immune suppression, angiogenesis, and treatment resistance, making it a critical target for therapy [64]. Efficient drug delivery remains a significant challenge in tumour nanomedicine, as many systems are trapped in the tumour ECM rather than penetrating deeper [65]. Computational and *in vitro* studies [63] show that nanoparticle diffusion decreases with increasing ECM density and stiffness, yet the experimental literature on this topic is limited. Our QDs-based approach offers a promising solution for studying diffusion in such microenvironments. Additionally, live-tissue dyes [66] (e.g., 5-ALA, Fluorescein Sodium, ICG) can complement our method for straightforward diffusion measurements in live tissue. In addition, the simplicity of QD surface modification makes them ideal for improving drug delivery. As dyes and reporters used in our imaging techniques are bath-applied, it is, therefore, possible to explore *ex vivo* in surgically resected human brain tissues or post-mortem samples collected with a reasonable time lag to validate the penetrability of genuine AD plaques or tumours by reporters with dimensions comparable to therapeutics.

In conclusion, our study provides new insights into the factors that influence the penetration of exogenous molecules into Aβ plaques themselves and Aβ plaques-bearing cortex in experimental AD. By elucidating the dynamics of the ECS in and around plaques, the findings advance our understanding of this devastating neurodegenerative disorder.

## 4. Experimental section/Methods

### Animals

Experiments were performed following the European Union directive (2010/63/EU) on protecting animals used for scientific purposes. They were approved by the Ethical Committee of Bordeaux University (CE50, France) and the Ministry of Education and Research under license number APAFIS #32540-2021072016125086 v11. The study is reported following ARRIVE guidelines (https://arriveguidelines.org).

8-month-old female APPswe695/ PS1ΔE9, termed APP/PS1 (Stock number: 005864), obtained from Jackson Laboratory (Bar Harbor, ME, USA), and their wild-type (WT) littermates (C57BL6/J) were used. Plaque size diversity (Figure 4) was investigated at the revision stage using a batch of animals from the same source (4 APP/PS1 animals, female aged 7/10 months). Briefly, the APP/PS1 mice express a chimeric mouse/human amyloid precursor protein APPswe (mouse APP695 harbouring a human Aβ domain and mutations K595N and M596L linked to a Swedish familial AD) and a human presenilin 1 mutated in familial AD (PS1ΔE9; deletion of exon 9). These bigenic mice were created by co-injection of both transgenes allowing for a co-segregation of the transgenes as a single locus [67]. Mice were generated in our animal facility from double-transgenic APP/PS1 males mated with C57BL/6 J females. Transgenic mice (APP/PS1) and age-matched non-transgenic littermates (WT) were allowed free access to food and water and maintained in a 12 h dark-light cycle. Mice were genotyped and systematically genotyped after each experiment (Transcriptomics Platform, Neurocentre Magendie, Bordeaux, France). In all experiments, females were used regardless of their oestrous cycle.

### In vivo TUSHI 2-photon microscopy

Animals were injected with buprenorphine (0.1 mg/kg) before the surgery for pain relief. Surgery was done under isoflurane anaesthesia. A round craniotomy (∼4 mm in diameter) left the dura mater intact, and was made above the somatosensory cortex. The dye (Alexa Fluor™ 488, carboxylic acid, Invitrogen, 3-4 µl with a concentration of 50 mM) was injected into the lateral ventricle on the side of the craniotomy at coordinates: M/L-1.1, A/P – 0.5, D/V – 2.3 at a rate of 500nl/min using a motorised syringe pump. The craniotomy was covered with a glass coverslip (#1 thickness, diameter 4 mm) and sealed with glue and dental cement (Superbond C&B). After surgery, mice were anaesthetized with ketamine/xylazine (100/10 mg/kg) and placed on a heated blanket under the objective of a custom-built 2-photon microscope, based on a commercial upright research microscope (BX51WI, Olympus, Hamburg, Germany) equipped with a 60X silicone oil objective (UPLSAPO, NA 1.3 Olympus) mounted on a z-axial nano-positioner (Pifoc 725.2CD, Physik Instrumente). 2-photon excitation was achieved using a femtosecond mode-locked fibre laser (Alcor 920, Spark lasers) delivering <100 fs pulses at a wavelength of 920 nm and a repetition rate of 80 MHz. Laser power was adjusted via a Pockels cell (302 RM, Conoptics) to up to 30 mW of power after the objective depending on imaging depth, and the pixel dwell time was 100 μs. The epi-fluorescence signal was descanned and detected by an avalanche photodiode (SPCM-AQRH-14-FC, Excelitas). Signal detection and hardware control were performed with the Imspector scanning software (Abberior Instruments) via a data acquisition card (PCIe-6259, National Instruments).

### In vivo QD tracking by 2-photon microscopy

Surgery was performed under isoflurane anaesthesia immediately before imaging. Animals were injected with buprenorphine (0.1 mg/kg) before the surgery for pain relief. A round craniotomy (1.2 mm in diameter) was made above the somatosensory cortex to expose the brain’s surface. Mice were placed on a heated blanket under the objective, and the head was fixed with a SGM-4 Head Holder for Mice (Narishige). 100 µL of Qdot™ 655 (Q11422 MP, Invitrogen) solution (1mM in aCSF) was placed over the brain surface and left for 5 minutes to diffuse into the tissue. The surface was then rinsed using physiological serum. Imaging was performed an Olympus BX61WI microscope (Olympus, Tokyo, Japan) equipped with a Nikon Apo LWD 25X water-immersion objective (NA 1.10). For 2-photon excitation, a Chameleon Vision 2 laser (Coherent, Santa Clara, CA, USA) was employed. The wavelength was set at 840 nm. Laser power modulation was controlled by a Pockels Cell (Conoptics, Danbury, CT, USA) in conjunction with a half-wave plate. Imaging was performed using the galvo scanner pathway, controlled by MESc software (Femtonics, Budapest, Hungary). Fluorescence signals were detected using a non-descanned GaAsP photomultiplier tubes (PMTs) equipped with red emission filters for signal collection. Acquisitions were performed between 20 and 40 micrometres depth. Videos were cropped based on intensity profiles using Python to eliminate the breathing effect and frames were 3-frame rolled averaged to improve signal. Analysis was performed with PALMtracer as described below.

### Acute brain slice preparation for ex vivo imaging

Mice were euthanised by cervical dislocation. Brains were swiftly extracted, and coronal sections (300-μm thick) were prepared in a VT1200S vibratome (Leica) in ice-cold NMDG solution and left to recover in NMDG solution for at least 20 min at room temperature. NMDG in mM: 93 NMDG, 2.5 KCl, 1.2 NaH_2_PO_4_·2H_2_0, 20 HEPES, 25 glucose, 30 NaHCO_3_, 10 MgSO_4_, 0.5 CaCl_2_. 1 Sodium Pyruvate and 12 N-acetylcysteine were added just before the experiment. pH was adjusted to 7.3-7.4 using HCl. Measured Osm 290-300. Slices were then transferred to room temperature aCSF (gassed with 95% O_2_, 5% CO_2_) with the following composition (in mM) 140 NaCl, 2.5 KCl, 1.2 NaH_2_PO_4_·2H_2_O, 26 NaHCO 3, 10 glucose, 10 HEPES, 2 MgSO_4_ and 2 CaCl_2_. Sodium Pyruvate and 12 N-acetylcysteine were added just before the experiment. pH was adjusted to 7.3-7.4 using HCl. The osmolarity was 300-310 mOsm.

### Ex vivo TUSHI 2-photon microscopy

Acute brain slices were imaged using a commercial 2-photon microscope (Prairie Technologies). For imaging of Calcein (200 µM, bath applied in aCSF composition: 126 NaCl, 3.5 KCl, 25 NaHCO_3_, 12 glucose, 1.2 NaH_2_PO_4_•2H_2_0, 1.3 MgCl_2_•6H_2_0 and 2 CaCl_2_•2H_2_0. Sodium Pyruvate and 12 N-Acetylcysteine, pH 7.3-7.4. The osmolarity was 300-310 mOsm). The wavelength of the 2-photon laser (Ti:sapphire, Chameleon Ultra II, Coherent) was tuned to 810 nm. Images were acquired using a 40X water immersion objective with an NA of 1.0 (Plan-Apochromat, Zeiss). Laser power was between 10 to 25 mW after the objective. The non-descanned fluorescence signal was collected by a PMT detector. Images were acquired with a pixel size of 144 nm over a 295x295 µm^2^ FOV (field of view) and pixel dwell-times between 15-25 μs. Image acquisition was controlled by the Prairie View software.

### TUSHI image processing and data analysis

TUSHI images were visualised and analysed using ImageJ. We manually selected ROIs using the “free-shape” drawing tool and calculated surface area of amyloid and crown (Figure 2a). The average total radius and average radius of amyloid (*r*) were calculated from the surface area (*A*) of amyloid + ring and amyloid, respectively, approximated by a circle from the equation *A* = *πr*^2^. The thickness of the ring was calculated as the difference of the total radius and radius of amyloid. All data in the text are represented as median and IQR.

### QD characterisation

Because of their proven bio-compatibility, we used commercial photoluminescent Qdot™ 655 (Q11422 MP, Invitrogen) [12]. These particles are passivated by a PEG corona, further decorated with F(ab’)2 IgG (H+L, goat anti-rabbit) fragments to confer antibody-like diffuse behaviour to the particles. Scanning transmission electron microscopy (STEM, Talos F200S G2 – Thermo Fisher Scientific Inc.) combined with a cationic negative staining protocol was used to image the nanocrystals’ dry size and the surrounding organic material (Supplementary Figure 3). Briefly, QDs were diluted in HEPES buffer (150 mM NaCl, 50 mM HEPES; pH 7.5) to a final concentration of 100 nM. A solution of uranyl acetate (UA) in diH_2_O (1.5 % w/v) was freshly prepared, protected from light, and filtered with a 0.22 µm cutoff. A 10 µL drop of QDs suspension was loaded on carbon grids (1 min., CF150-Cu-50 – Delta Microscopies), serial washes in diH_2_O (×3) followed and UA negative staining was performed (1 min.). A single wash in diH_2_O was finally done after staining (*note*: between each step, the liquid was removed through absorption by capillarity with filter paper). QDs on grids were imaged at accelerating voltages of 200 keV and visualised with a 4K×4K camera (One View – Gatan). The hydrodynamic size of QDs was studied by dynamic light scattering (DLS, Zetasizer µV – Malvern Panalytical). Nanoparticles were diluted in HEPES buffer to a final concentration of 50 nM and 50 µL of suspension was placed in a disposable cuvette. This latter was loaded and equilibrated on the device for 120 s at 20 °C. Readings consisted of averaging 8 measurements of 20 s repeated 4 times, count rate and correlation function were monitored to select reliable recordings in the absence of thermal drifts, sample agglomeration, and precipitation, as suggested by the manufacturer. Particle size distribution was fitted with a lognormal function to extract geometric mean and standard deviation.

### QD incubation in acute brain slices

Acute slices were prepared as described above. After preparation and recovery, slices were then incubated for 40 min at 35 °C in 5 mL of QDs suspension (200 pM) in carbogenated aCSF (95% O_2_, 5% CO_2_, in mM): 126 NaCl, 3.5 KCl, 25 NaHCO_3_, 12 glucose, 1.2 NaH_2_PO_4_•2H_2_0, 1.3 MgCl_2_•6H_2_0 and 2 CaCl_2_•2H_2_0. Sodium Pyruvate and 12 N-Acetylcysteine, pH 7.3-7.4. Measured mOsm 300-310. Slices were rinsed for 8 mins in 5 mL of bubbled QD-free aCSF before imaging.

### QD time-lapse confocal microscopy

Samples were placed in a thermostatic recording chamber (35 °C – Life Imaging Services GmbH) mounted on an upright microscope (Eclipse Ni-E, Nikon) equipped with a confocal spinning disk unit (CSU-X1, Yokogawa). Amyloid plaques in APP/PS1 mice cortices were identified by their autofluorescence under 405 nm (180 mW, 5%) laser diode excitation (L4Cc, Oxxius), while QDs were excited at 488 nm (200 mW, 50%). Excitation light was separated from the emitted using a quad-band polychroic mirror (405/488/561-568/635-647, Chroma®); emitted light was further passed respectively on a 447/60 and 655/15 nm single-band bandpass filter (BrightLine®). During time-lapse particle tracking, slices were continuously perfused with gassed aCSF at a rate of 1 mL/min (35 °C). Cortical areas were identified under 10× magnification (CFI Plan Fluor, 0.45 N.A.), a 60× water dipping objective (CFI Apo NIR, 1 N.A.) was instead used to image senile plaques and QDs (> 15 µm depth). Particle diffusion was sampled at 28 Hz for 1 min using an EMCCD camera (Evolve® 512 – Teledyne Photometrics). The imaging system was controlled by an integrated acquisition software (NIS-Elements AR – Nikon). For plaque size diversity identification, recordings were performed as explained above, but keeping the precise location of each plaque. After several QDs recordings, ThioflavinS 0.01% was added for 10 minutes and then rinsed by aCSF perfusion to label amyloid fibres. A z-projection of maximal fluorescence intensities across 10 optical slices (Z-step = 1 μm) was made through the recordings of QDs. The plaque area was selected using the *Wand tracing Tool* after “isodata” thresholding. The selection was used to calculate circularity using the Fiji parameter “Shape Descriptor” (circularity = 4π x area /(perimeter) 2). Categorical classification was performed as described in [10]: Filamentous plaques had a circularity score of 0.00–0.14, and compact plaques had circularities greater than 0.28. Plaques with circularity scores between 0.15–0.28 were classified as having ‘intermediate’ phenotypes.

### QD localisation and trajectory analysis

Low-resolution image stacks of single emitters were segmented, particle position was super-localized and single molecule trajectories were reconnected with PALMtracer, a high-end software package (MetaMorph® – Molecular Devices) for the analysis of single-molecule dynamics. Briefly, a wavelet segmentation and a Gaussian fitting (6 pixels around maxima) are used for threshold-based localisation. At the same time, a simulated annealing algorithm is applied for trajectory reconstruction through successive images. As a result, the mean-square displacement (MSD) of moving particles as a function of time is obtained. Only trajectories described by more than 10 points were analysed. The instantaneous diffusion coefficient (D_inst_) was calculated for all trajectories through a linear fit of the first 4 points of the MSD curve. At the curve plateau, between 0.6 and 0.8 s, the mean square displacement average value was used as a proxy of the confinement area experienced by QD. To evaluate the extent of trapping within the plaques, the immobile fraction was obtained by counting the number of particles moving at velocities lower than 0.005 µm^2^/s over the total (only FOV where *Amyloid* ROI had at least one immobile QD were considered for immobile fraction analysis). Relative changes of studied parameters in the three (*Out*, *Ring* and *Amyloid*) anatomical compartments of the plaques were studied as average percentage variation between adjacent ROI in the same 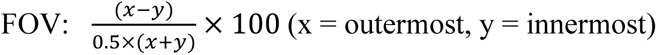.

### QD localisation accuracy

The localisation accuracy for QD tracking was measured with the SPT setup described above (see *QD time-lapse confocal microscopy*). About 50 frames were recorded to establish an immobile particle’s *x* and *y* location within the field of view. Particles with median velocities lower than 0.005 µm^2^/s were considered immobile [68]. The standard deviation (s.d.) of particle position in each frame was measured, and the averages between *x* and *y* s.d. were plotted. Instantaneous diffusion of immobile particles over time was also plotted to highlight the difference between QD immobilisation on a glass coverslip (which defines our SPT setup limit of localisation accuracy) and immobile QD detected within the living tissue (which concerns the actual experimental localisation). See Supplementary Figure 6

### SWCNTs preparation and characterisation

SWCNTs solution was prepared following the previously described methods [35] with some minor modifications. 1 mg HiPco synthesised CNTs (batch #189.7) was suspended in 0.5% w/v phospholipid-polyethylene glycol (PL-PEG) (#mPEG-DSPE-5000, Nanocs) in 10 mL MQ water. The dispersion was typically homogenised for 15 min at 8,000 rpm using a dispersing instrument (IKA T-10 Basic), and then further dispersed by tip sonication at 20 W (output power) for 10 min under the ice. Nanotube bundles and impurities were precipitated by centrifugation at 10621g for 60 min at room temperature. 70–80% of the supernatant of PL-PEG suspended SWCNTs was then collected and stored at 4 °C. The concentration of the CNTs stock solution was estimated by UV (EVOLUTION 220, Thermo Scientific). A range of SWCNTs concentrations have been used (1 – 10 mg/L). The length of the SWCNTs was measured by AFM. For this, typically, SWCNTs solution was drop-casted on freshly cleaved mica surfaces. Then, the substrate was gently rinsed with water, followed by overnight air drying (Supplementary Figure 11).

### SWCNT intracerebroventricular injection

6 μl of SWCNT solution (2-10 mg/mL) were injected into the right lateral ventricle of living mice by stereotactic surgery (coordinates from bregma: −0.5 mm anteroposterior, 1 mm mediolateral, −2.4 dorsoventral) as previously described [35]. 30 min before surgery, animals received subcutaneously received buprenorphine (0.1 mg/kg) as an analgesic and lurocaine (5 mg/kg) as a local anaesthetic. Surgery was performed under deep isoflurane anaesthesia. SWCNTs solution was injected with a 30-gauge Hamilton syringe coupled to a microinjection pump (World Precision Instruments) at a 1 μl/min flow rate. The needle was left in place for 12 min to avoid leakage, then slowly retracted, stopping for another 2 mins at -1.2 DV. Between 40 min-1 hour after an injection of the nanotubes, mice were euthanised by cervical dislocation.

### Ex-vivo SWCNT near-infrared microscopy

Acute slices were prepared as described above. After recovery, images were collected at 35 °C in a 3D-printed chamber with controlled temperature by a feedback system. Pre-warmed carbogen-bubbled aCSF was perfused throughout the chamber by a peristaltic pump. Slices were imaged for no more than 45 min. Imaging of moving SWCNTs in the ECS was performed on a customized upright epifluorescent microscope (Nikon). An 845 nm laser diode (QSI), polarised circularly with a quarter-wave plate (Thorlabs), was used to excite the (6,5) SWCNTs at a phonon sideband. Emission light was collected through a 900nm long-pass filter (Chroma) using a 25x/1.10NA water-dipping objective (Nikon). Images were acquired with an InGaAs camera (C-RED 2 – First-Light Imaging) at 33 ms exposure time. A 1.5x zoom magnification was applied to match the camera pixel size with the microscope point-spread function width. Before recording, transmission white light was used to check the position in the entire slice and determine the brain region to be imaged. To avoid non-physiological data acquisition, the first 10 μm of tissue was always discarded to exclude the first cell layer, which was potentially damaged during slicing.

### SWCNT SPT analysis

The analysis was performed as previously described with minor modifications [35]. Briefly, individual SWCNTs were super-localized with a 2D Gaussian fit, and coordinates were linked to reconstruct individual trajectories. Local diffusion and local ECS dimensions were then extracted as previously described in [36] upon drift correction using non-mobile particles in the field of view. For each trajectory, the instantaneous mean square displacement (MSD_inst_) analysis was used to estimate the instantaneous diffusion coefficient D_inst_. MSD values were calculated over a sliding window of 10 frames and linear fits were then applied to the first 3 time points to retrieve values of D_inst_. Immobile SWCNTs, characterised by the plateau shape of their global MSD, were excluded from the analysis. To estimate local ECS dimensions, the shape of the local area explored by individual SWCNTs along their trajectory was analysed using a 6-point time window [36]. This time window was chosen at maximum confinement (i.e. when the shape of the local area is maximally distorted by local ECS dimension as compared to expected in unconfined environments) as defined by the eccentricity ratio of the ellipse formed by the SWCNT trajectory (Supplementary figure 12).

### SWCNT localisation accuracy

The precision of our measurements is critically influenced by both the number of photons detected and the level of background noise. We analysed three representative trajectories of diffusing single-walled carbon nanotubes (SWCNTs) and extracted the localisation precision using ThunderSTORM [69], as shown in Supplementary Figure 13.

Assuming a median localisation precision of 30 nm, we can estimate the error on the measured instantaneous diffusion coefficients owing to this precision. From the equation of the mean squared displacement (MSD), we find that the error on the instantaneous diffusion coefficient can be approximated as:

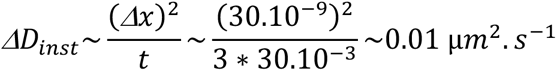

Where *Δx* is the localization precision and t is the time window where *ΔD_inst_* Is measured. This error is in the order of ∼15% of the median value of D_inst_ found in this study.

### Immunostaining and fluorescence imaging

Brains were swiftly extracted and fixed in 4% PFA at 4 °C for 24h and rinsed in PBS. Then, 50 μm-thick vibratome coronal sections were collected and kept in Phosphate-buffered saline (PBS) at 4°C before immunostaining. For visualisation of the hyaluronan matrix, were treated with a streptavidin-biotin blocking kit (Vector Labs) for 20 min and incubated overnight with biotinylated-hyaluronic acid binding protein (HABP from bovine nasal cartilage, Merck-Millipore) diluted in blocking solution. Staining was revealed with Streptavidin-Atto647N (Sigma-Aldrich). Double or triple labelling was achieved by re-blocking samples with 4% normal goat serum and overnight incubation with primary antibodies for the following antigens: Iba1 (Rb, 019-19741, Wako), GFAP (Ms, MAB360, Merk Millipore). Staining was revealed with appropriate secondary antibodies conjugated with Alexa 488 or 594 (Thermo Fisher). Aggrecan was revealed by overnight incubation with Aggrecan Polyclonal Antibody (Rb,13880-1-AP, Thermo Fisher). PNNs were labelled with Wisteria Floribunda Lectin (WFA, WFL) conjugated with a green fluorophore (L32481, ThermoFisher) by incubation for 6h. Sections were mounted on #1.5 coverslips with VECTASHIELD® Antifade Mounting Medium and left to dry overnight in darkness. Confocal images were acquired in a Leica TCS SP8 microscope with a 63X Plan Apo CS objective with oil immersion, maintaining image acquisition settings (laser power, AOTF, detection parameters) between sessions. Image stacks: pixel size: 90 nm, z-step 0.5 μm. Images were treated and analysed with Fiji/ImageJ. Wide-field images (PNN imaging) were acquired in a BX 63 Olympus microscope with a mercury lamp, using a Dry UPLFLN 10X Objective (NA 0.3). Acquisition parameters were kept between acquisitions. Analysis was performed in Image/J Fiji. ROIs were drawn manually. Each value was obtained from 3 slices covering the sensory-motor cortex of mice.

### Expansion microscopy

Double labelling was achieved on 50 μm-thick vibratome coronal sections (HABP from bovine nasal cartilage, Merck-Millipore) and β-Amyloid (Biolegend) (adapted from [70]). Staining was revealed with appropriate secondary antibodies conjugated with Alexa 488 or 568. Samples were incubated overnight at RT with the succinimidyl ester of 6- ((acryloyl)amino) hexanoic acid (acryloyl-X, SE; abbreviated AcX; Life Technologies),1/100 in PBS. Gelation was achieved by incubation for 30 min at 4°C in momomer solution (1XPBS,2M NaCl, 8.625% (w/w) sodium acrylate, 2.5% (w/w) acrylamide, 0.15 (w/w) N,N’-methylenebisacrylamide) was mixed and ammonium persulfate (APS) initiator and tetramethylethylenediamine (TEMED) accelerator were added up to 0.2% (w/w) each and inhibitor 4-hydroxy-2,2,6,6-tetramethylpiperidin-1-oxyl (4-hydroxy-TEMPO) up to 0.01% (w/w) before using. Slides were placed in a chamber at 37°C for 2 hours -digestion and expansion: gelled samples were then incubated in proteinase K 8 units/mL in digestion buffer (50mM Tris (pH8), 1mM EDTA, 0.5% Triton X-100, 0.8M guanidine HCL) for at least 12 hours at RT. Digested gels were next placed in excess volumes of doubly de-ionized water for 2 hours to expand. Images were imaged in water in a Leica TCS SP5 with an objective 25x water immersion objective (NA 0.95), maintaining image acquisition settings (laser power, AOTF, detection parameters) between sessions. Image stacks: pixel size: 110.2 nm, z-step optimized by software. Images were treated with Fiji/ImageJ.

### Quantitative real-time PCR (qPCR)

Total RNA was isolated using the Quick-RNA™ FFPE Kit (ZYMO research). RNA was processed and analysed following the MIQE guidelines [71]. cDNA was synthesised from 0.5 μg of total RNA by using qSript XLT cDNA SuperMix (Quanta Biosciences). QPCR was performed using a LightCycler® 480 Real-Time PCR System (Roche). QPCR reactions were duplicated for each sample, using transcript-specific primers, cDNA (4 ng) and LightCycler 480 SYBR Green I Master (Roche) in a final volume of 10 μl. The PCR data were exported and analysed in an informatics tool (Gene Expression Analysis Software Environment) developed at the NeuroCentre Magendie. The RefFinder method was used [72] to determine the reference gene. Relative expression analysis was normalised against two reference genes, and the succinate dehydrogenase complex subunit (Sdha) and Elongation factor 1-alpha 1 (Eef1a1) genes were used. The comparative (2^-ΔΔCT^) method [73]calculated the relative expression level. Primer sequences are reported in Supplementary Table 1.

### Statistical analysis

TUSHI: Data was analysed in ImageJ Fiji and MS Excel; statistical analysis was done in Graph Pad Prism 9. SPT data (including QD and SWCNT) was analysed Graph Pad Prism 9. Matrix complexity data was analysed using Graph Pad Prism 9. QPCR analysis and statistics were performed using the *GEASE* tool (Developed by Neurocentre Magendie, Bordeaux, France).

## Acknowledgements

We thank Sebastien Marais, Melina Petrel, Sabrina Lacomme, Marie-Laure Arotcarena and Claire Mazzocco for their help in setting up the experimental conditions. We also thank Nathan Benac, Morgane Meras, Flavia Simões, Remi Kinet and Morgane Darricau for helpful discussions. We thank Aude Panatier for providing supplemental animals needed at revision stage. Electron, expansion, and in vivo SPT microscopy were done in the Bordeaux Imaging Centre, a service unit of the CNRS-INSERM and Bordeaux University, a member of the national infrastructure France BioImaging supported by the French National Research Agency (ANR-10-INBS-04). We thank the Cell Biology Facility (IINS) for cellular tool productions and the genomic platform at Neurocentre Magendie for qPCR analysis.

## Author contributions

Conceptualization: JEP, IC, YD, VN, EB

Methodology: JEP, IC, YD, VN, EB

Investigation: JEP, IC, YD, SN, QG, ED, TLL

Resources: TA

Supervision: LG, LC, VN, EB

Writing—original draft: JEP, YD, IC, VN, EB

Writing—review & editing: All authors contributed and revised the final version

## Data and materials availability

The source data for all graphs and charts are available at https://doi.org/10.5281/zenodo.15056592 (DOI)

## Code availability

Customs scripts and software code generated for this paper are available from the corresponding authors upon reasonable request.

## Supporting Information

Supporting Information is available from the Wiley Online Library or from the author.

## Supplementary Materials for

**This PDF file includes:**

Figs. S1 to S13

Supplementary Table 1

**Other Supplementary Materials for this manuscript include the following:**

Movies S1 to S9

## Supplementary Figures

**Supplementary Figure 1.**
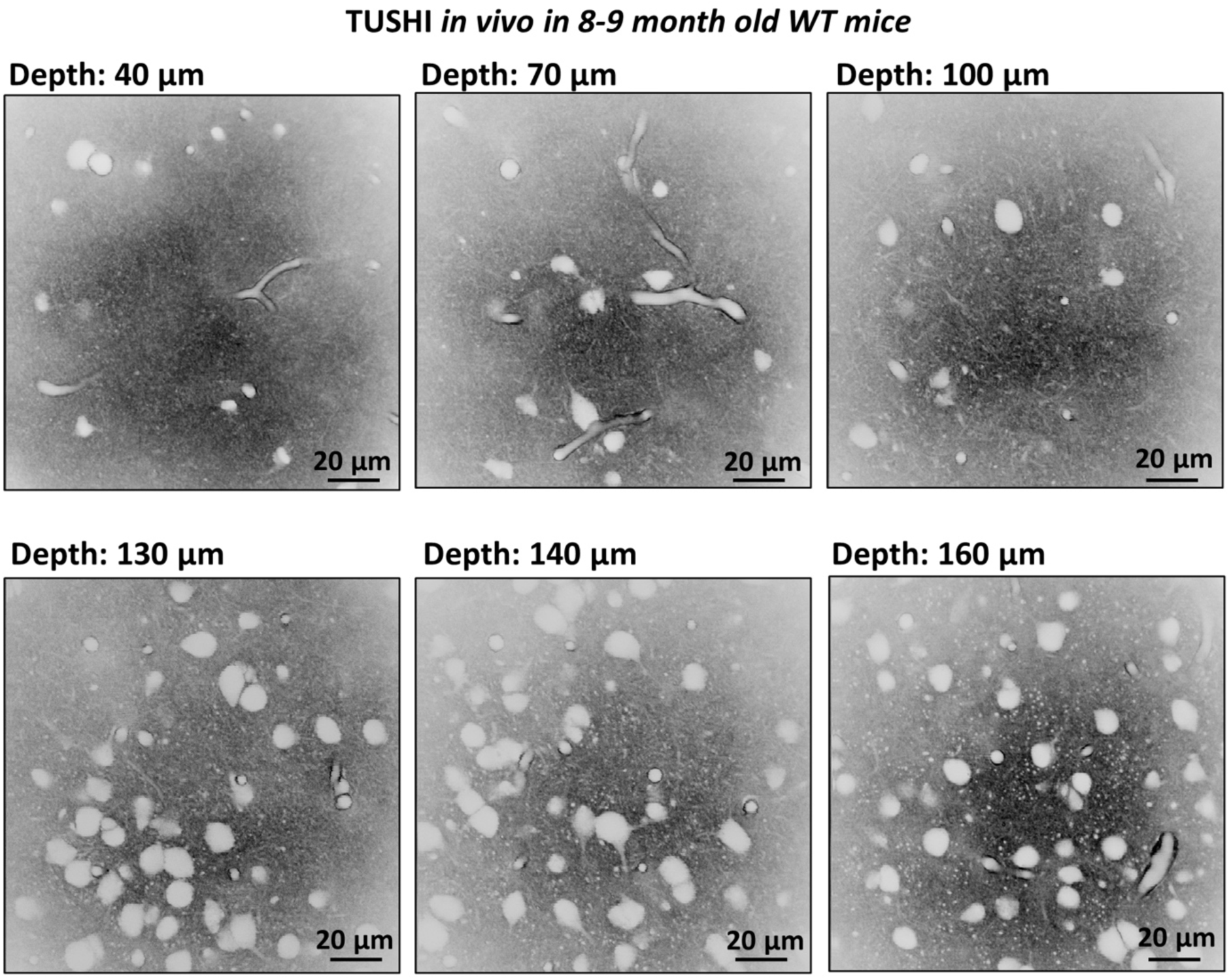
Visualisation of the cortex using TUSHI in 9 9-month-old WT mice in *in vivo* condition.

**Supplementary Figure 2.**
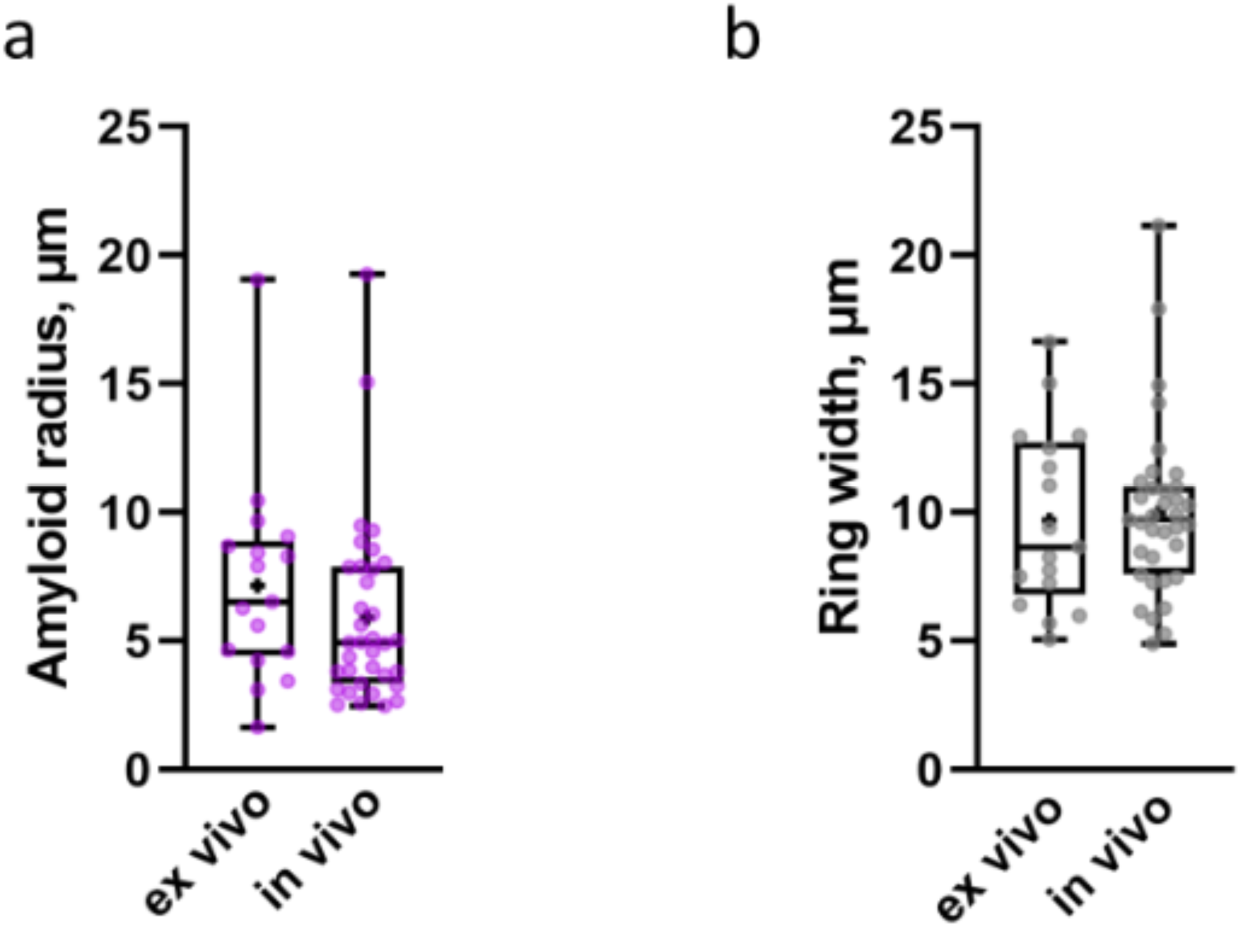
Measurements of the size of amyloid core (a) and the ring (b) using TUSHI in *ex vivo* and *in vivo* conditions.

**Supplementary Figure 3.**
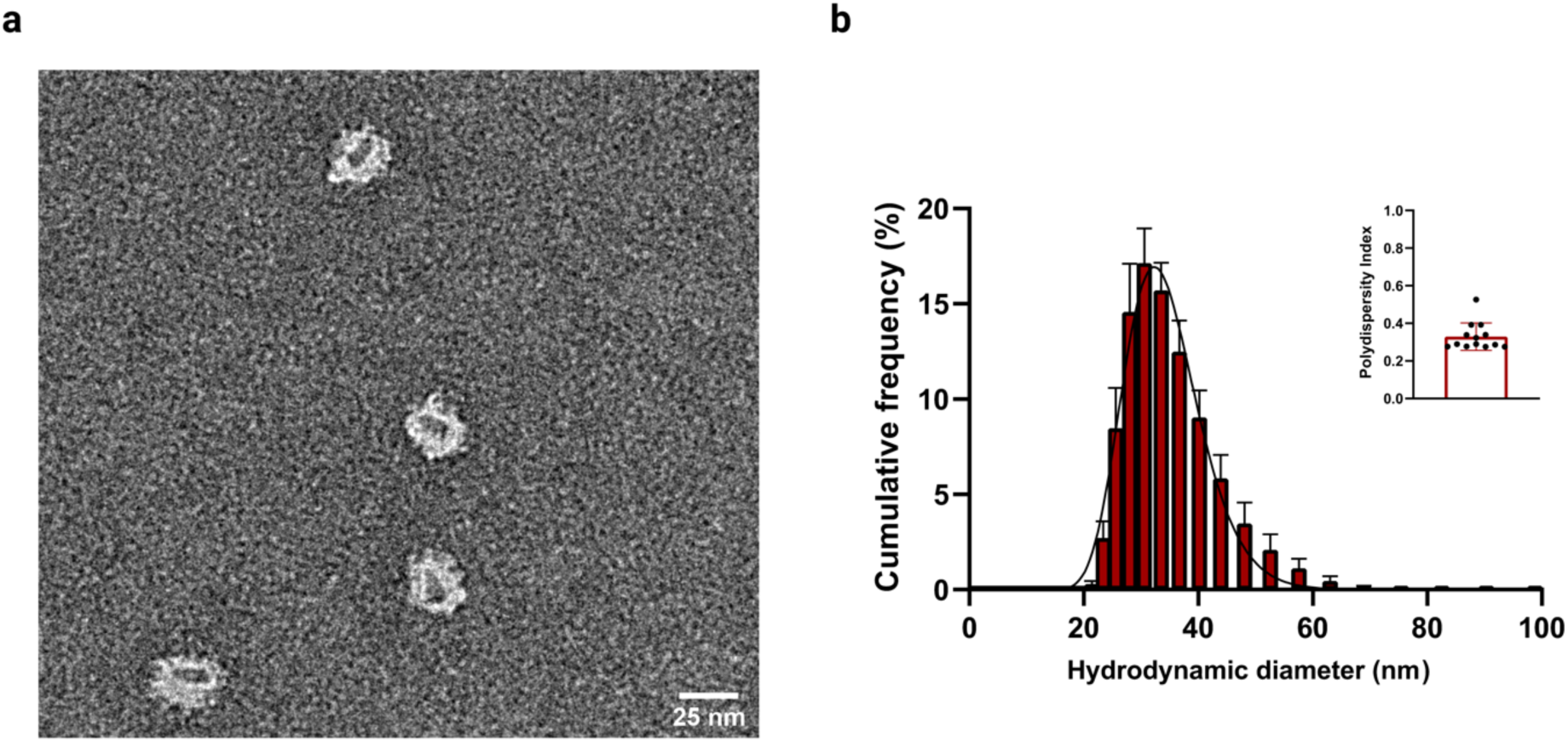
Charactezitacion of Quantum Dots. (a) Transmission electron microscopy (TEM) of negatively stained QDs provides the dry size of QD. The metallic, electron-dense, core (dark) is distinguishable from the surrounding polymer-protein (bright) shell (25 nm). (b) Log-normal fitted distribution of particles hydrodynamic diameters (33.6 ± 1.2 nm) obtained by dynamic light scattering (DLS) in HEPES-buffered saline (pH 7.5). Top right inset reports the low polydispersity index (0.29, IQR = 0.28-0.53) for size distributions from individual replicates.

**Supplementary Figure 4.**
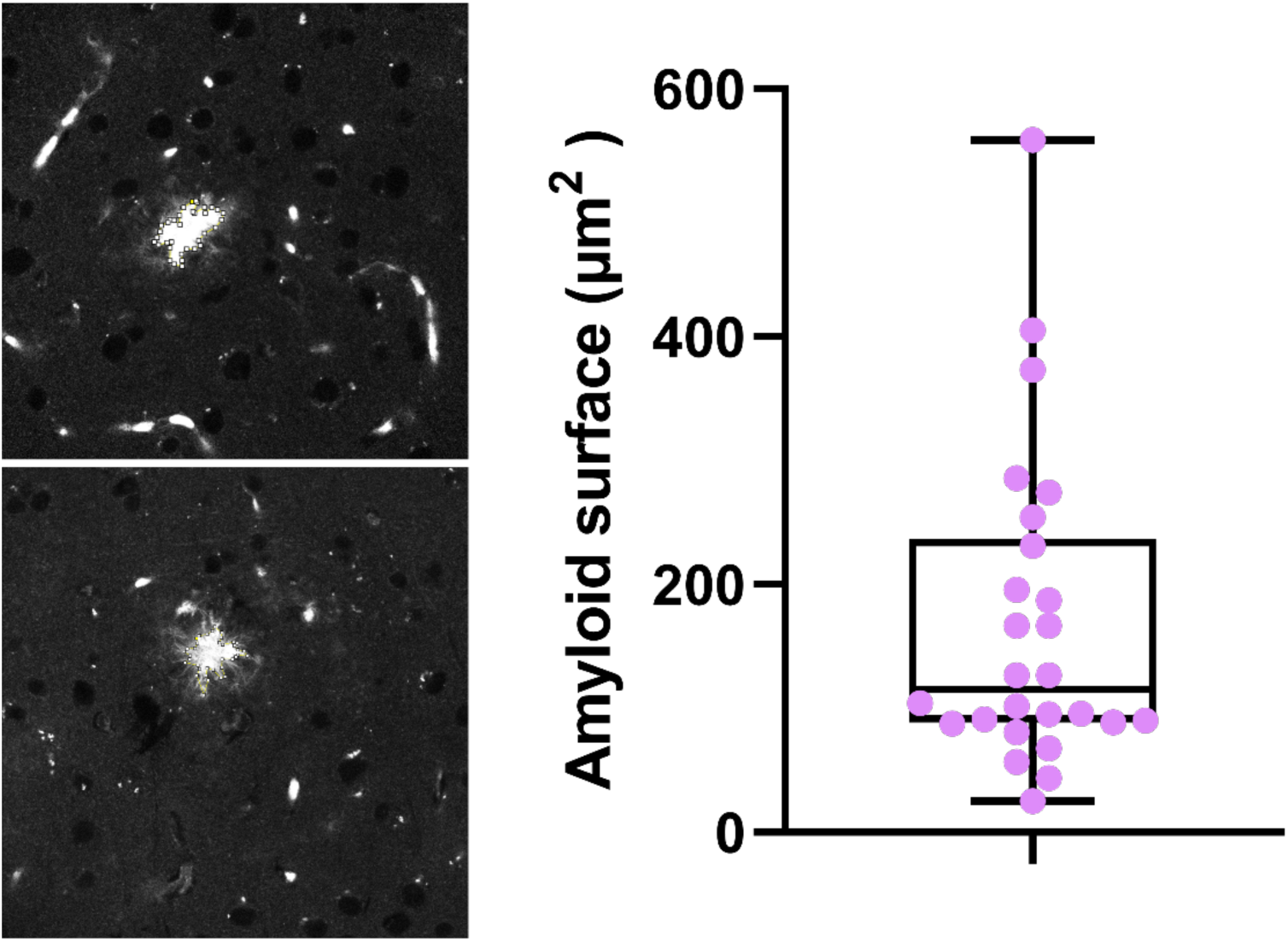
Amyloid surface quantified by autofluorescence. (In μm^2^: Median = 115, IQ25 = 89, IQ75 = 237, 26 plaques from n=4 mice).

**Supplementary Figure 5.**
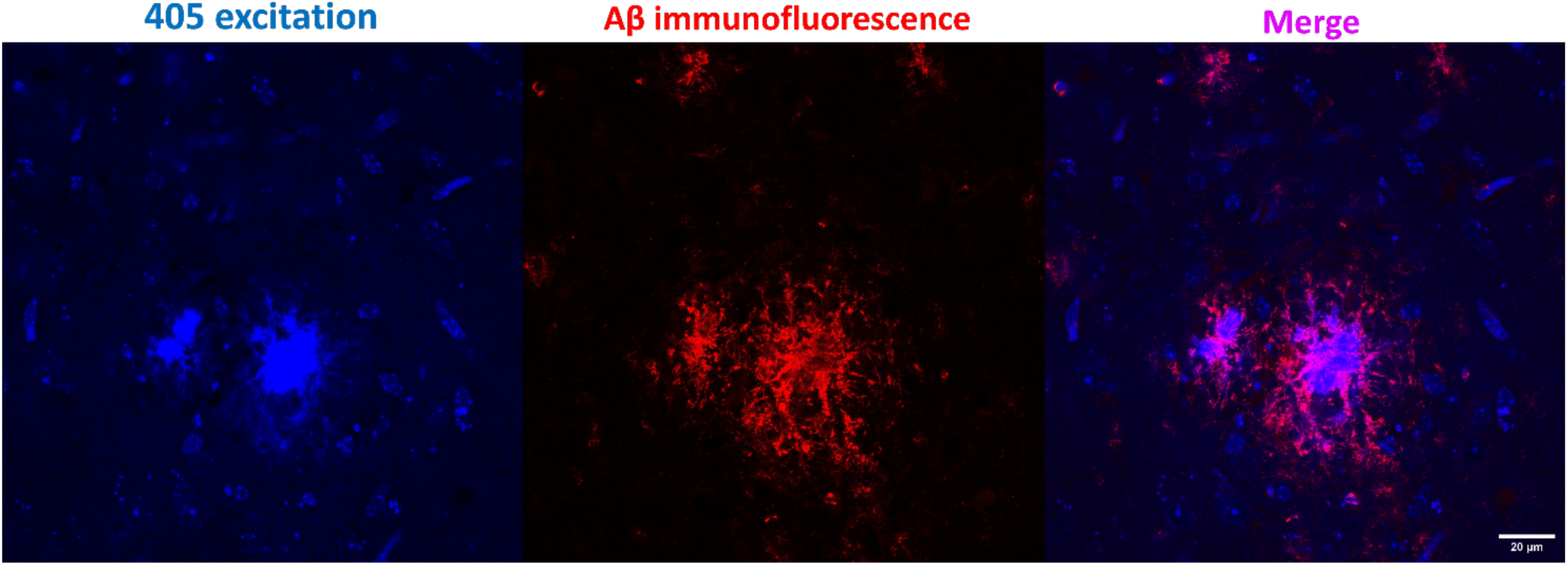
Blue autofluorescence upon 405 excitation colocalises with immunofluorescence signal against Anti-β-Amyloid, 1-16 (6E10, Bio legend).

**Supplementary figure 6.**
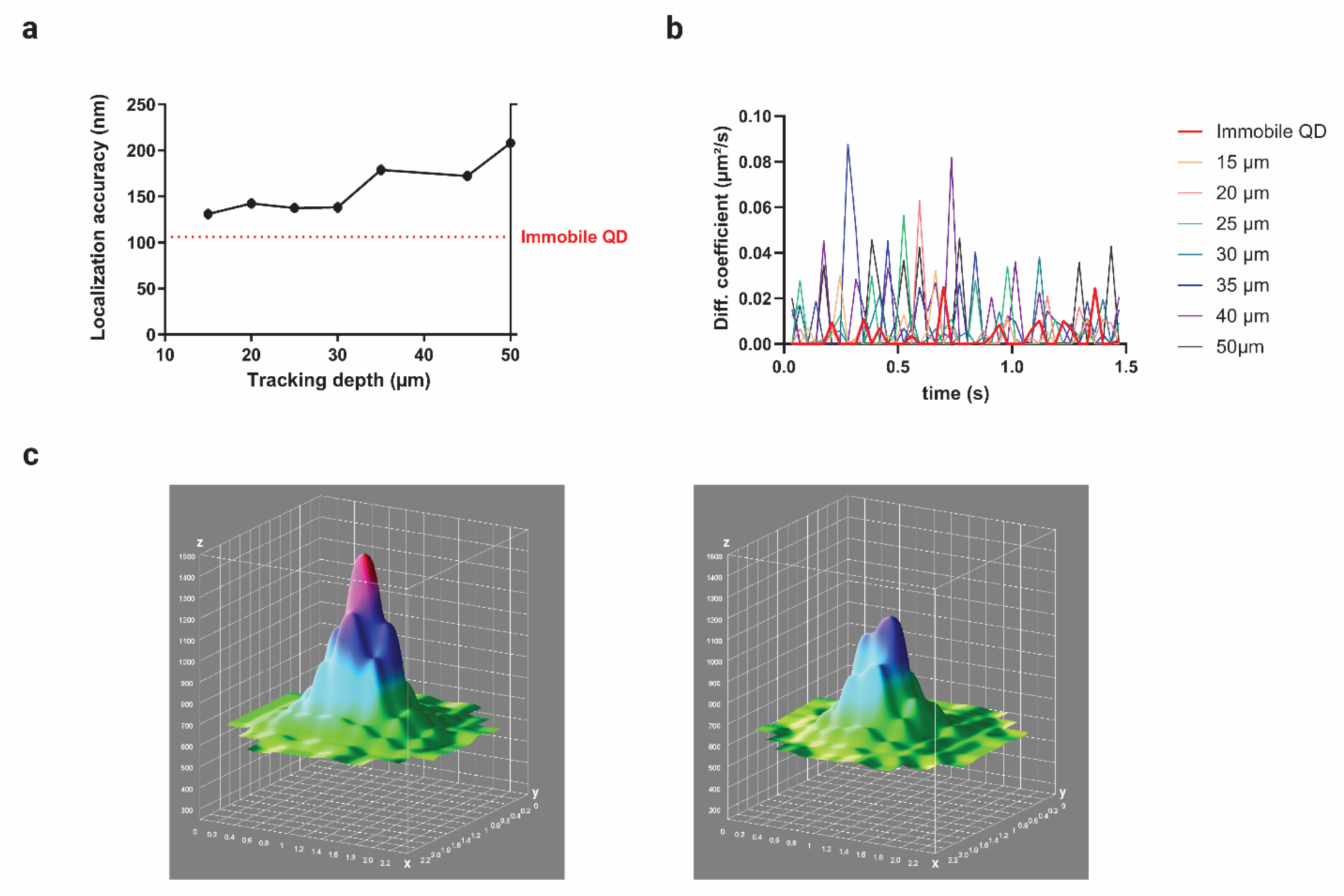
Localisation accuracy of QDs tracked on upright spinning disk microscope. (a) The super-resolved localisation limit of the SPT setup is obtained from immobile quantum dots (dashed red line) adsorbed and immobilised on a glass slide; virtually immobile particles within living cortical slices were also super-localized at different depths (black line) to highlight the interdependence between the two parameters (which is also a consequence of collected photon number). Measured values agree with previous results obtained with a comparable setup (see ref. (Biermann et al., 2014). (b) Instantaneous diffusion of a truly immobile (thick red line) particle on a glass slide, and virtually immobile particles (coloured lines), at various depths within the tissue, is also given to explain the difference between perfect immobility and immobile regime in the tissue; the former displaying lower speed fluctuations compared to immobile particles within the tissue, which can experience instantaneous nanometric displacement as a consequence of tracking in a living system. (c) Two 3D surface plots of immobile QDs are shown for comparable localisation accuracies obtained at 20 µm (left plot) and at 30 µm (right plot) depths within the tissue. The greater signal at 20 µm is shown to justify limiting the study to such depth; tracking at increased depths comes with a lower detection of photons, thus affecting the signal over the background noise.

**Supplementary Figure 7.**
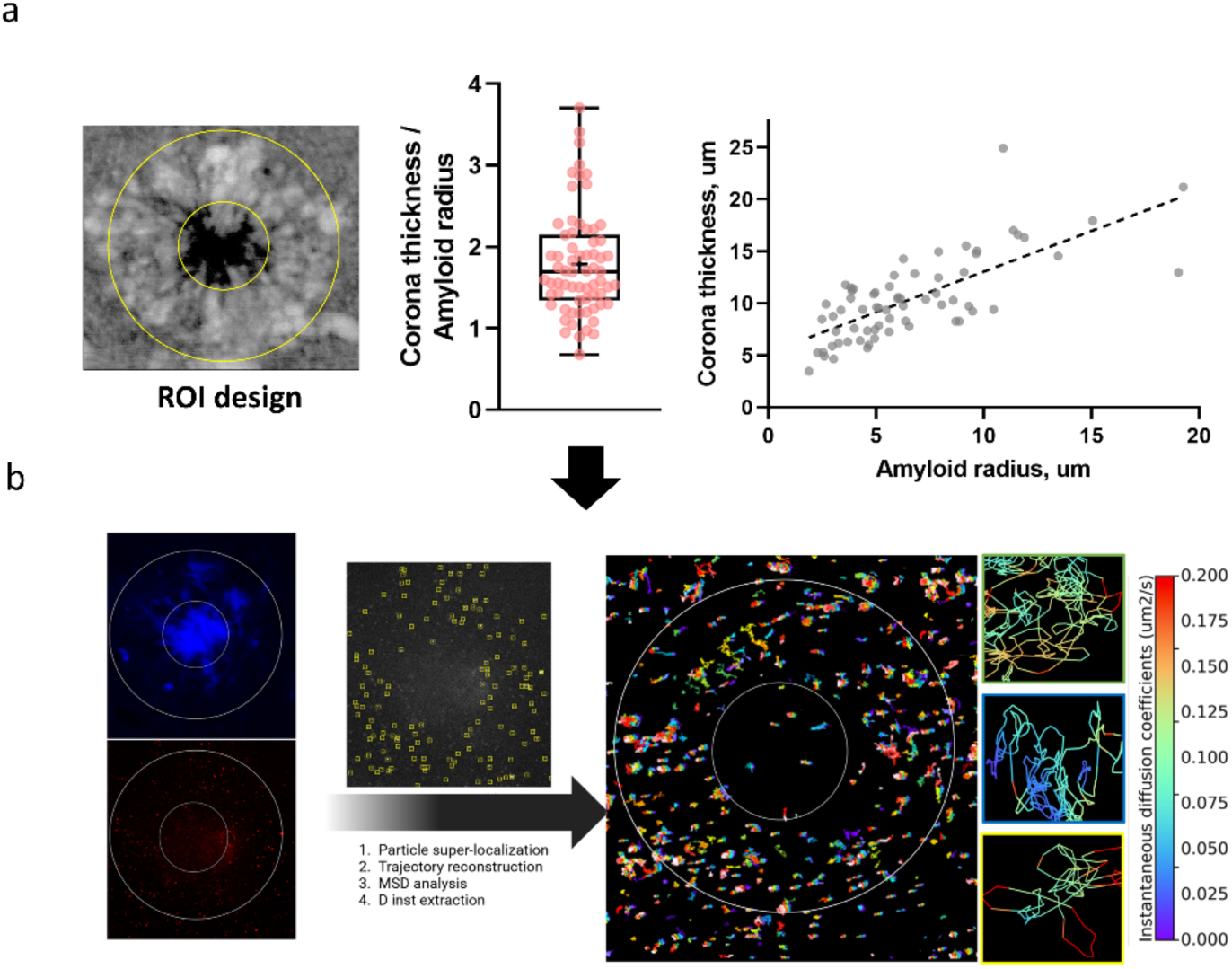
ROI design from TUSHI to QDs SPT analysis. (a) After conducting shadow imaging analysis, we observed that the region of interest (ROI) associated with the ring could be represented as concentric to the amyloid core, exhibiting a median width 1.82 times larger than the amyloid ratio. This relationship demonstrated a strong correlation (r = 0.73, p < 0.0001), as illustrated in Figure 2. (b) ROIs for QDs signal were drawn based on the proportions described by shadow imaging. A first ROI was drawn using the threshold blue autofluorescence signal and a second following the mentioned proportions. Analysis of QDs diffusion was performed with *PALMtracer* (MetaMorph® – Molecular Devices). Briefly, the analysis involved 1) particle super localisation, 2) trajectory reconstruction and MSD analysis, and 3) D_inst_ value extraction. Each of the values was ascribed to each of the ROIs designed beforehand. The result gauges the building of a map with the trajectories of particles.

**Supplementary Figure 8.**
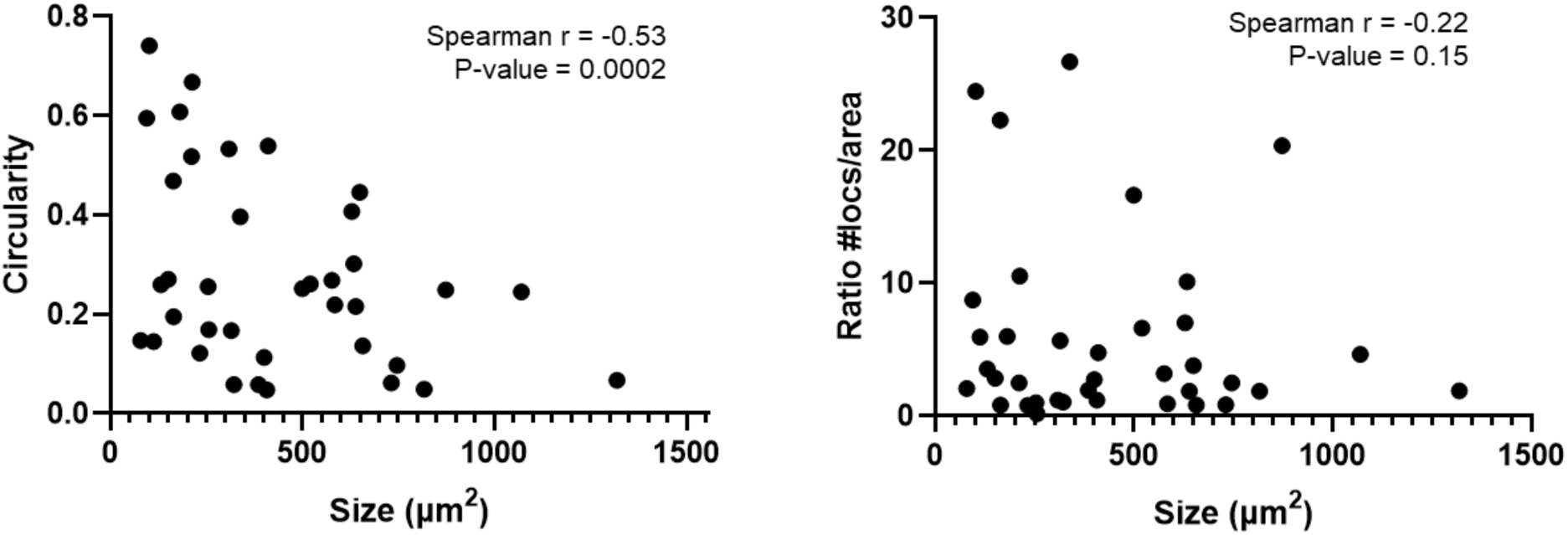
There is a strong negative correlation between circularity and sizes (Spearman r =-0.53, p = 0.0002). However, there is no significant correlation between size and ratio locs/area (Spearman r = -0.22 p = 0.15).

**Supplementary Figure 9.**
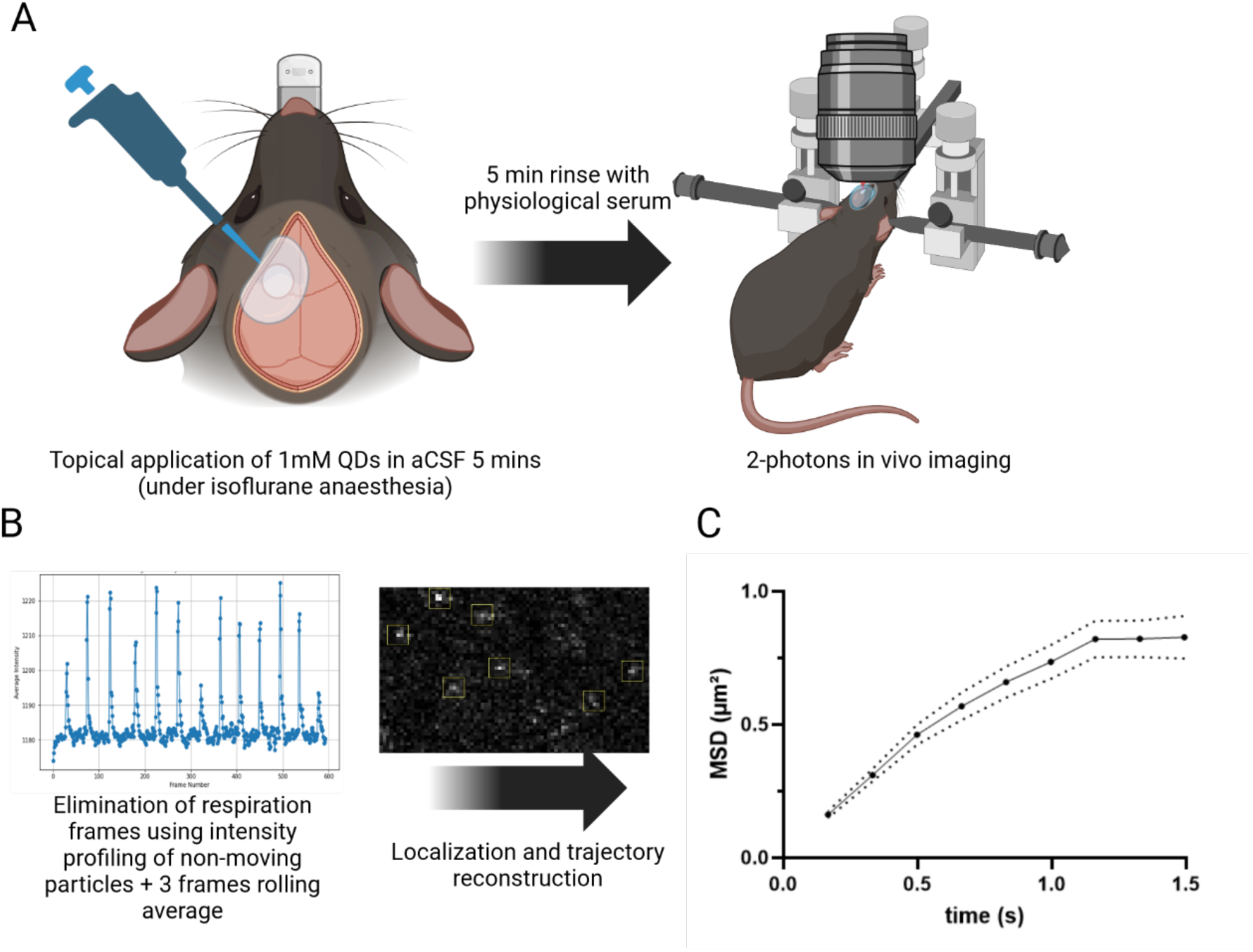
In vivo QDs SPT tracking experimental protocol. (a) A round craniotomy (1.2 mm in diameter), was made above the somatosensory cortex to expose the surface of brain. 100µL of QDs solution (1mM in aCSF) was placed over the brain surface and left for 5 minutes to diffuse into the tissue. The surface was then rinsed using physiological serum. The mice were then imaged in a 2-photon setup. (b) The analysis involved the elimination of respiration frames. A 3-point rolling average was then applied to improve signal. Localization and trajectory reconstruction were performed in MetaMorph**®**. (c) The total MSD shows the characteristic shape of restricted diffusion, the same found in *ex vivo*. A total of 1048 MSDs from 8 recordings (n = 2 mice) were computed in the MSD graph.

**Supplementary Figure 10.**
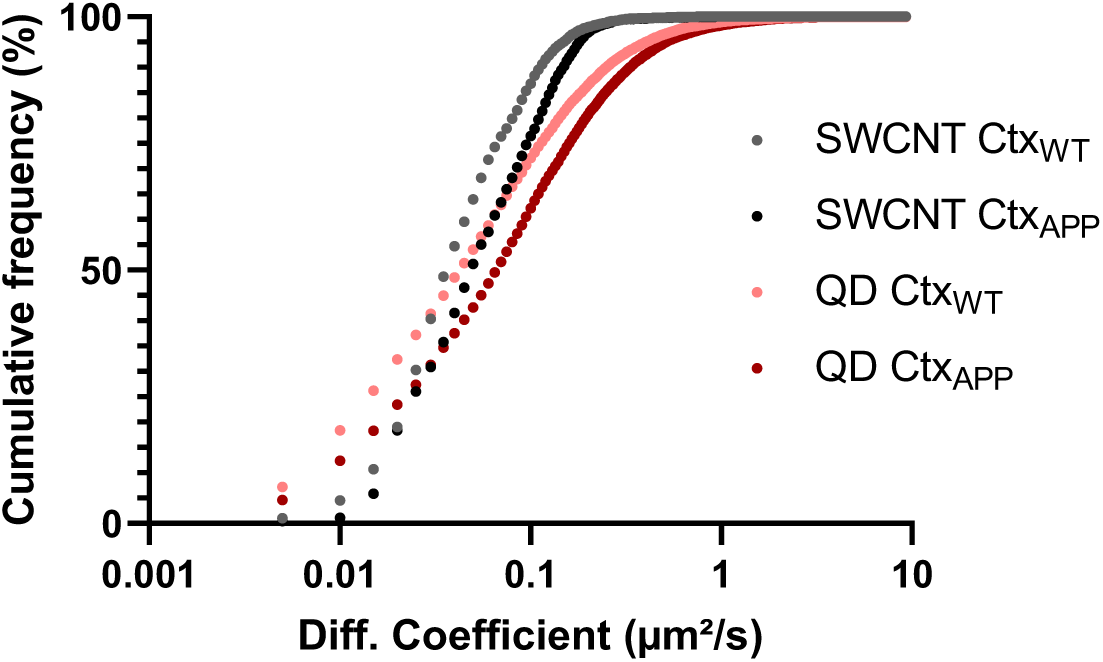
Supplementary Figure 10. Comparative of QDs and SWCNTs diffusion coefficients in WT and APPPS1 tissue. Quantum Dots (QDs) and Single-Walled Carbon Nanotubes (SWCNTs) exhibit a comparable pattern of enhanced diffusion in the amyloid brain vs Wild-type. However, their respective instantaneous diffusion coefficients vary, likely reflecting the influence of the probe. (QDs: WT Median = 0.045, IQR = 0.017-0.115, APPPS1 Median = 0.069, IQR = 0.024- 0.16; SWCNTS: WT Median = 0.038, IQR = 0.025-0.069, APPPS1 Median = 0.051, IQR = 0.027-0.98).

**Supplementary Figure 11.**
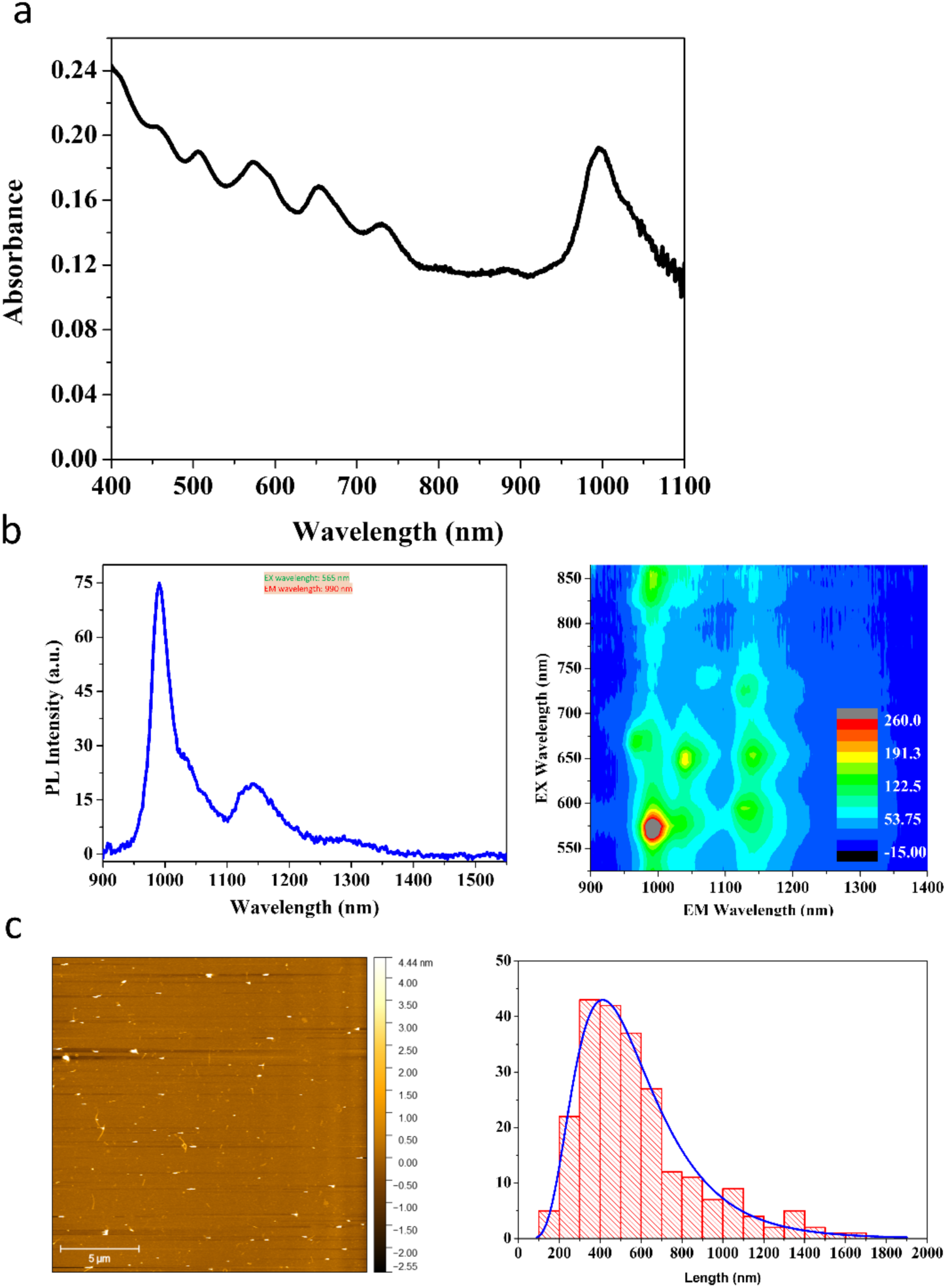
SWCNTs spectroscopic characterisation. (a) The absorption spectrum of the HiPco synthesised CNTs suspended in 0.5% w/v PLPEG. Measured in EVOLUTION 220, Thermo Scientific (b) PL spectra of the PLPEG suspended SWCNTs with their corresponding 2D excitation-emission PL map (NanoLog, HORIBA). (c) AFM image of the PLPEG suspended SWCNTs with their length distribution. Mean length = 575 ± 285 nm (Median length = 510).

**Supplementary Figure 12.**
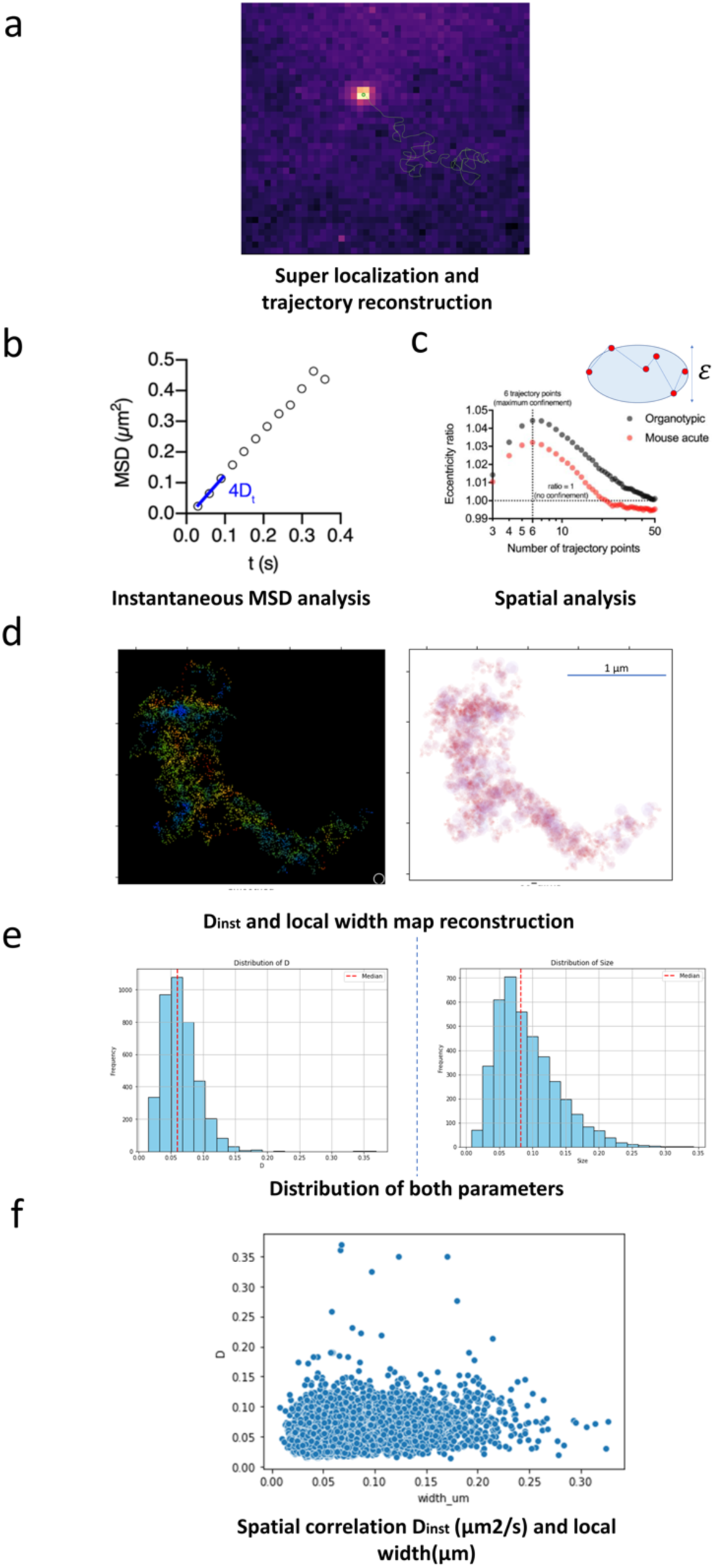
Schematics of SWCNTs analysis. (a) Individual SWCNTs were super-localised with a 2D Gaussian fit, and coordinates were linked to reconstruct individual trajectories. (b) For each trajectory, iterative analysis of the instantaneous mean square displacement (MSD_inst_) was used to estimate the instantaneous diffusion coefficient D_inst_. MSD values were calculated over a sliding window of 10 frames, and linear fits were applied to the first 3 time points to retrieve D_inst_’s values. (c) To estimate local ECS dimensions, the shape of the local area explored by individual SWCNTs along their trajectory was analysed using a 6-point time window. This time window was chosen at maximum confinement (i.e., when the local area’s shape is maximally distorted by local ECS dimension compared to expected in unconfined environments) as defined by the eccentricity ratio of the ellipse formed by the SWCNT trajectory. Each SWCNTs trajectory rendered a (d) nanoscale map of D_inst_ and local width, a (e) distribution of D_inst_ and local width values and (f) spatial correlation information between both types of data.

**Supplementary Figure 13.**
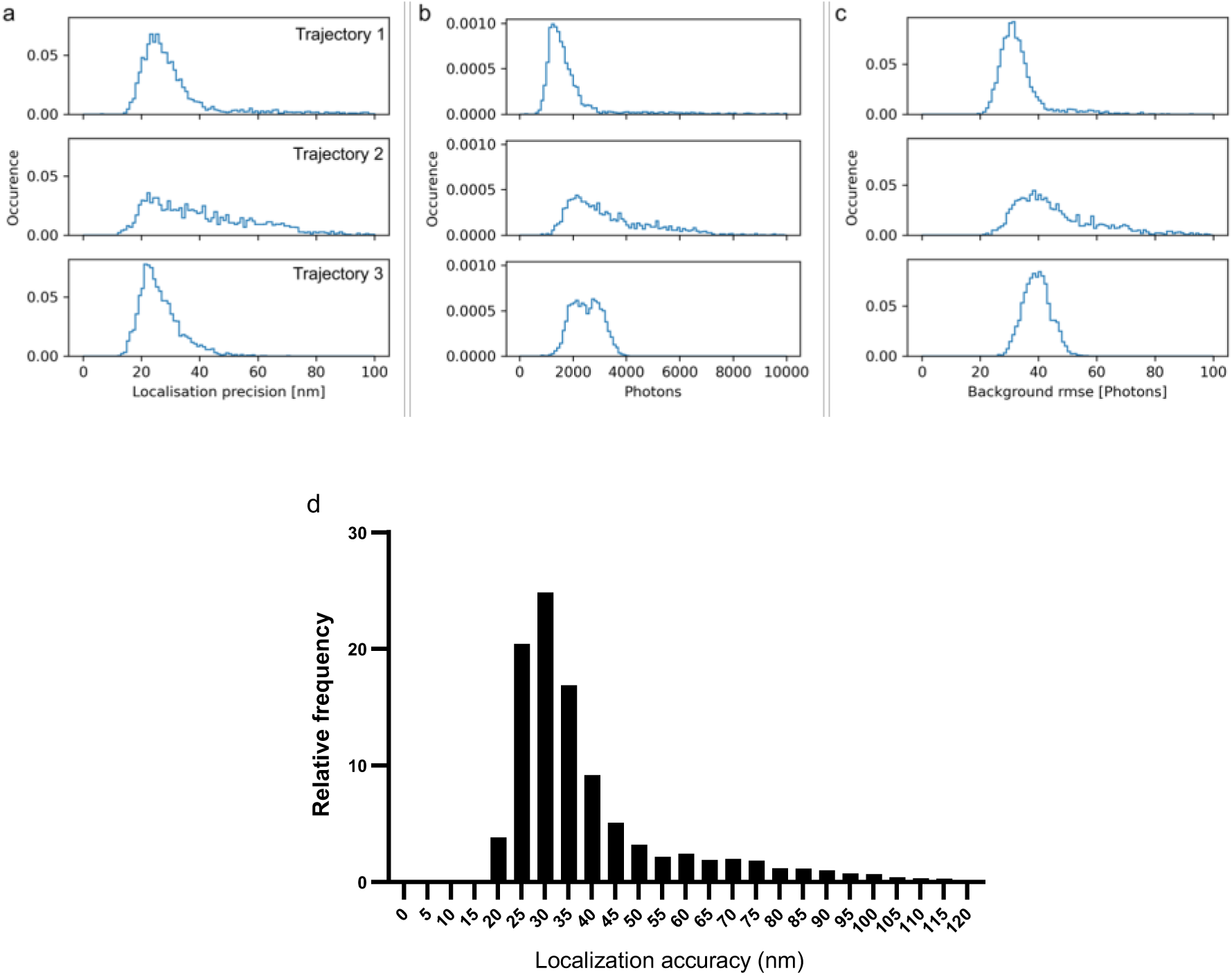
Localization precission of SWCNTs. (a) Localization precision, (b) number of photons, and (c) background noise, computed from the three representative trajectories of diffusing SWCNTs in the brain extracellular space (ECS). (d) Histogram showing localization precisions extracted from 3 previous trajecotries. Median = 32,8 IQR = 27.6-42.5.

**Supplementary Table 1.**
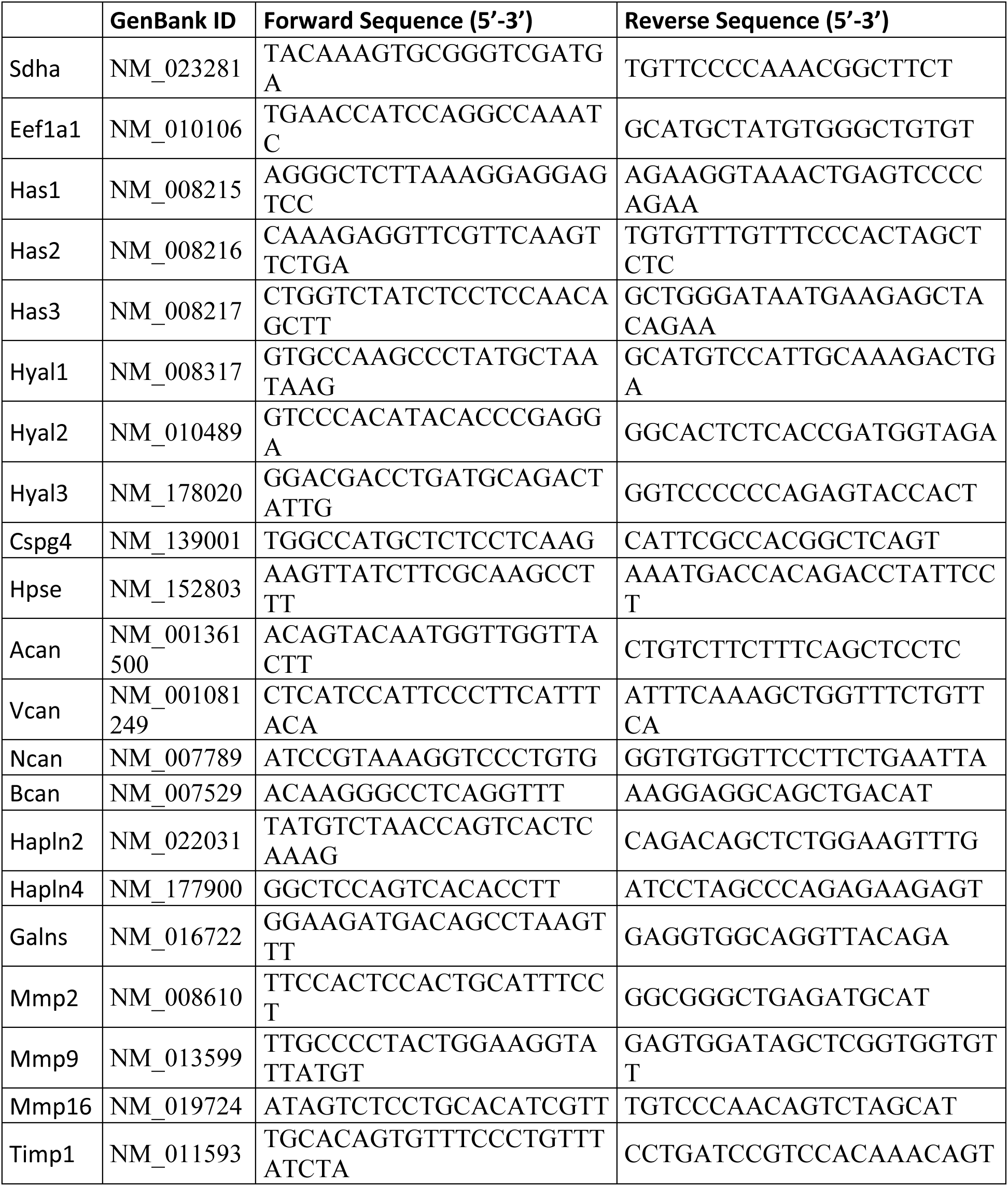
qPCR primer sequences.

**Movies**

**Supplementary video 1.** *Ex vivo* (acute slice) imaging of amyloid plaques using TUSHI in the cortex of an APP/PS1 mouse.

**Supplementary video 2.** *In vivo* imaging of amyloid plaques and ring using TUSHI in the cortex of an APP/PS1 mouse.

**Supplementary video 3.** *In vivo* imaging of amyloid plaques and ring using TUSHI in the cortex of an APP/PS1 mouse.

**Supplementary video 4.** *In vivo* imaging of the cortex using TUSHI in a WT mouse.

**Supplementary video 5.** *In vivo* imaging of the cortex using TUSHI in a WT mouse.

**Supplementary video 6.** *Ex vivo* (acute slice) imaging of QDs diffusing around a cortical amyloid plaque in an acute brain slice of an APP/PS1 mouse (Depth = 20μM).

**Supplementary video 7**. *Ex vivo* (acute slice) imaging of QDs diffusing in the cortex of a WT mouse (Depth = 20μM).

**Supplementary video 8.** *In vivo* tracking of QDs in the cortex using 2p microscopy in a WT mouse.

**Supplementary video 9.** *In vivo* tracking of QDs in the cortex using 2p microscopy in a WT mouse after breathing frames elimination and denoise (rolling average).

Biermann, B., et al., 2014. Imaging of molecular surface dynamics in brain slices using single-particle tracking. Nature communications. 5, 3024.

