## Supplementary Figure 1 for "Decoding Amyloid Plaque Penetrability: Exploring Extracellular Space and Rheology in Plaque-rich Cortex"

**This PDF file includes:**

Figs. S1 to S13
Supplementary Table 1

**Other Supplementary Materials for this manuscript include the following:**

Movies S1 to S9

**Supplementary Figures:**

**Fig S1**

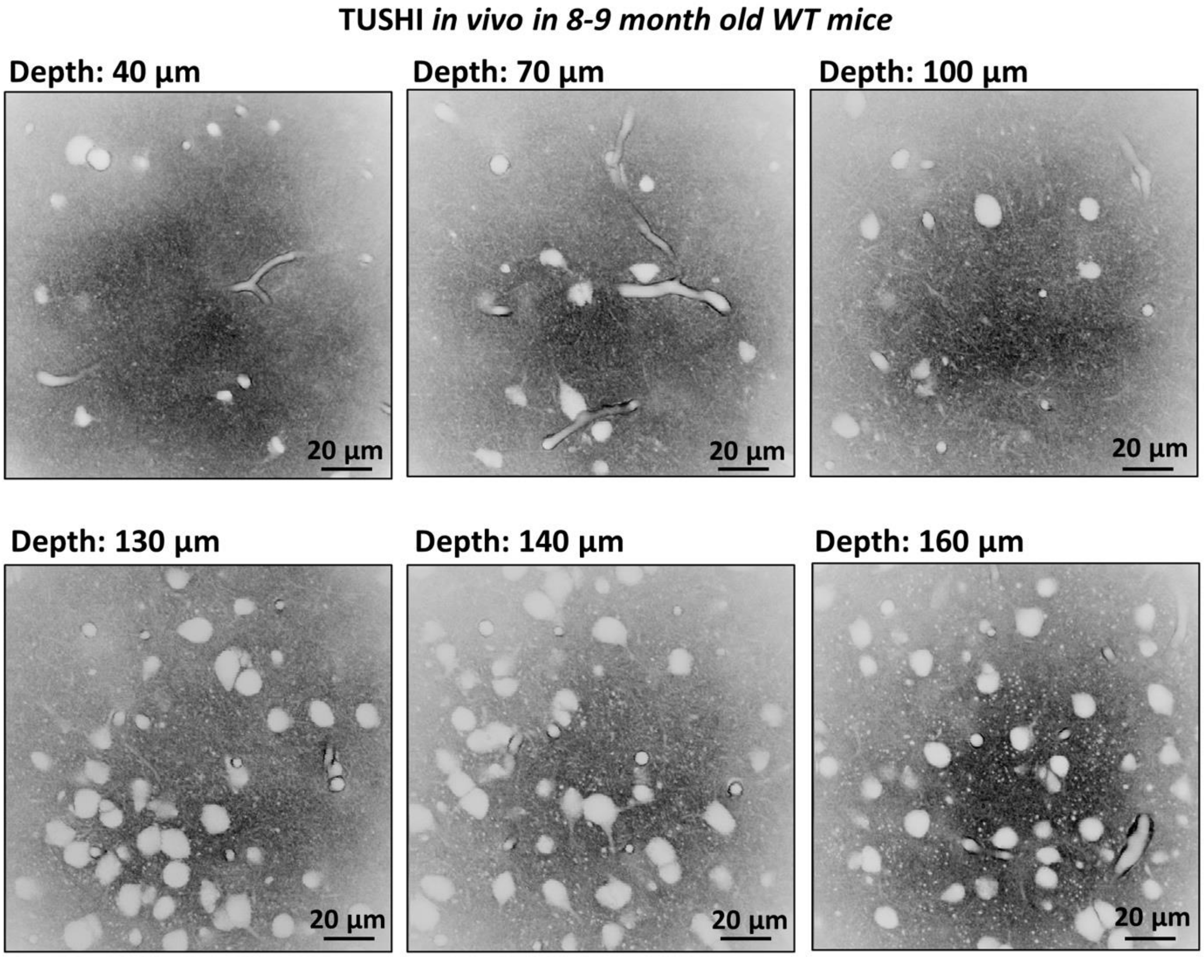


**Supplementary Figure 1**. Visualisation of the cortex using TUSHI in 9 9-month-old WT mice in *in vivo* condition.

**Fig S2**

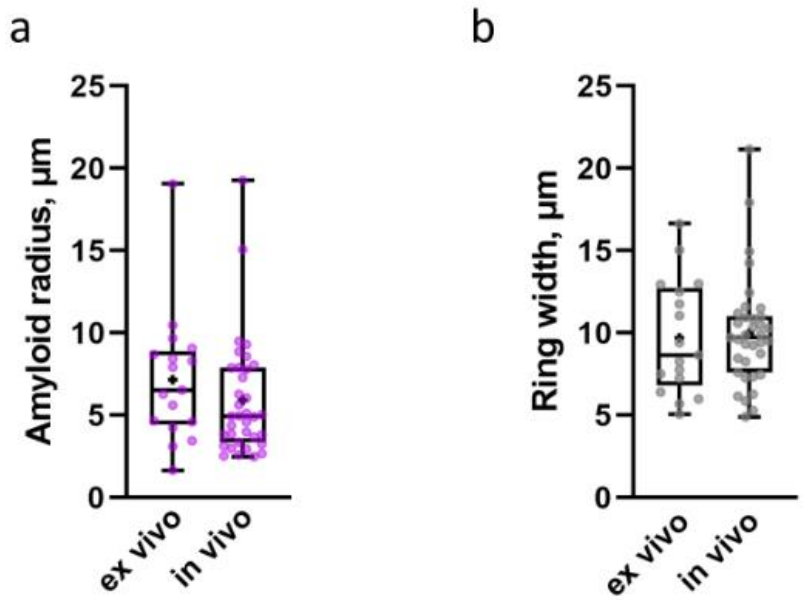


**Supplementary Figure 2.** Measurements of the size of amyloid core (a) and the ring (b) using TUSHI in *ex vivo* and *in vivo* conditions.

**Fig S3**

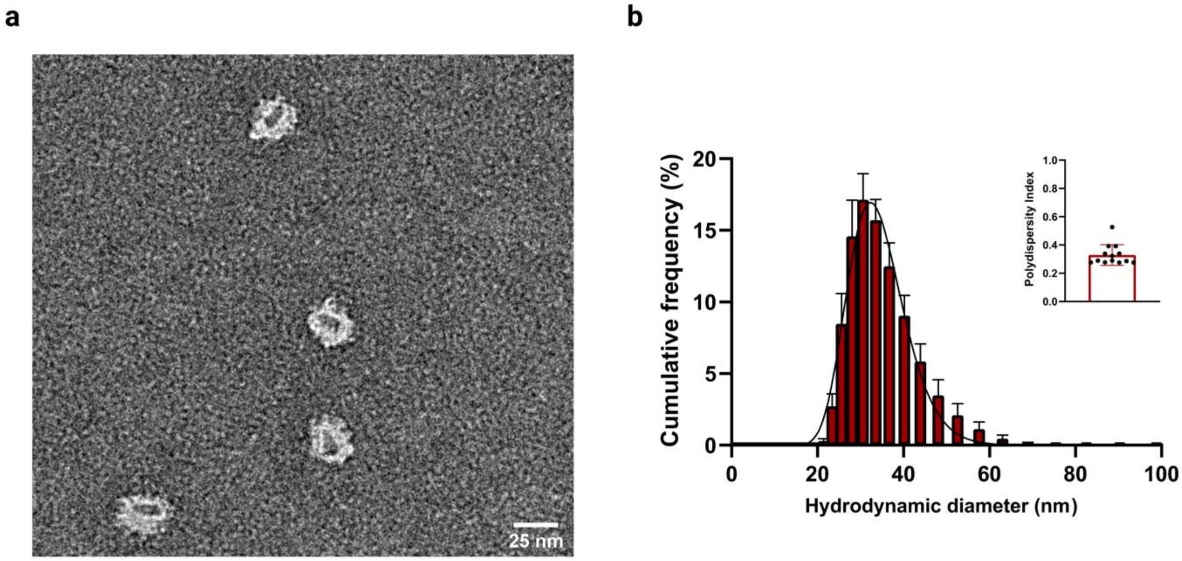


**Supplementary Figure 3. (**a) Transmission electron microscopy (TEM) of negatively stained QDs provides the dry size of QD. The metallic, electron-dense, core (dark) is distinguishable from the surrounding polymer-protein (bright) shell (25 nm). (b) Log-normal fitted distribution of particles hydrodynamic diameters (33.6 ± 1.2 nm) obtained by dynamic light scattering (DLS) in HEPES-buffered saline (pH 7.5). Top right inset reports the low polydispersity index (0.29, IQR = 0.28-0.53) for size distributions from individual replicates.

**Fig S4**

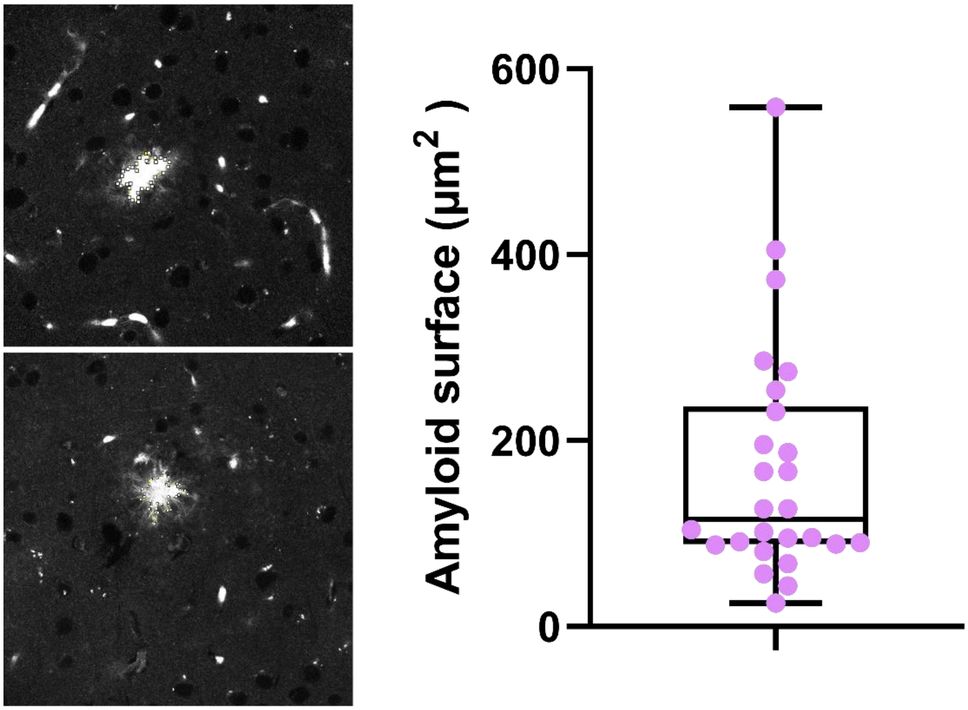


**Supplementary Figure 4.** Amyloid surface quantified by autofluorescence. (In μm$^2$: Median = 115, IQ25 = 89, IQ75 = 237, 26 plaques from n=4 mice).

**Fig S5**

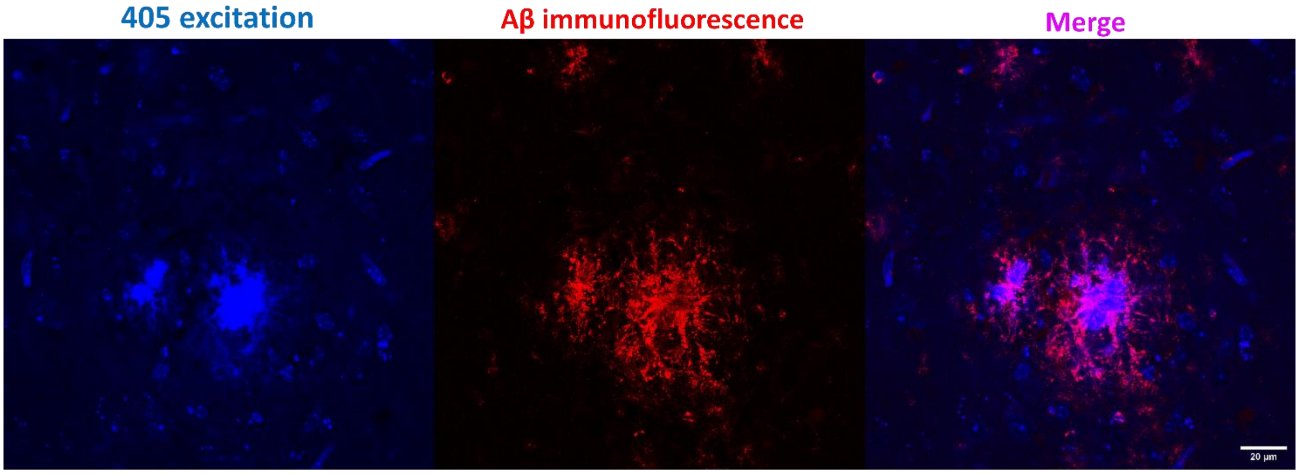


**Supplementary Figure 5**. Blue autofluorescence upon 405 excitation colocalises with immunofluorescence signal against Anti-β-Amyloid, 1-16 (6E10, Bio legend).

**Fig S6**

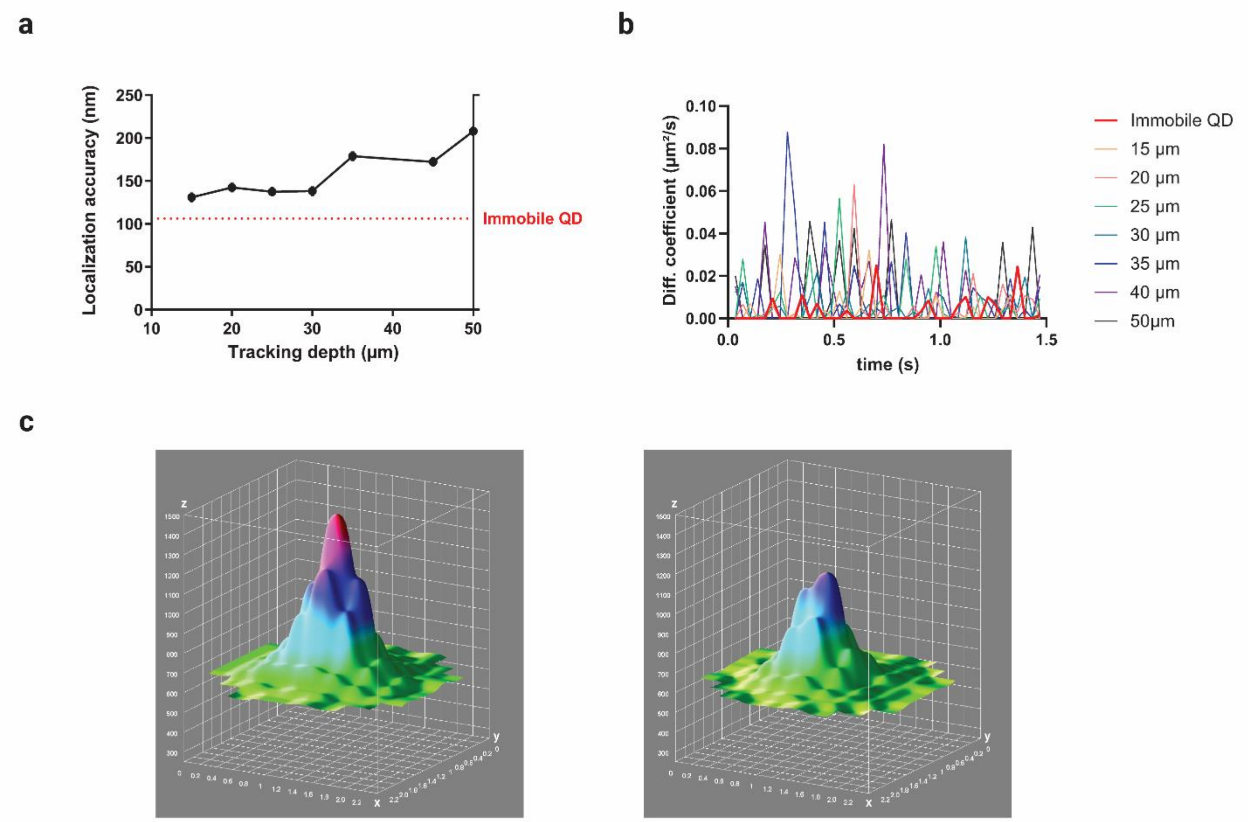


**Supplementary figure 6.** Localisation accuracy of QDs tracked on upright spinning disk microscope. (a) The super-resolved localisation limit of the SPT setup is obtained from immobile quantum dots (dashed red line) adsorbed and immobilised on a glass slide; virtually immobile particles within living cortical slices were also super-localized at different depths (black line) to highlight the interdependence between the two parameters (which is also a consequence of collected photon number). Measured values agree with previous results obtained with a comparable setup (see ref. (Biermann et al., 2014). (b) Instantaneous diffusion of a truly immobile (thick red line) particle on a glass slide, and virtually immobile particles (coloured lines), at various depths within the tissue, is also given to explain the difference between perfect immobility and immobile regime in the tissue; the former displaying lower speed fluctuations compared to immobile particles within the tissue, which can experience instantaneous nanometric displacement as a consequence of tracking in a living system. (c) Two 3D surface plots of immobile QDs are shown for comparable localisation accuracies obtained at 20 µm (left plot) and at 30 µm (right plot) depths within the tissue. The greater signal at 20 µm is shown to justify limiting the study to such depth; tracking at increased depths comes with a lower detection of photons, thus affecting the signal over the background noise.

**Fig S7**

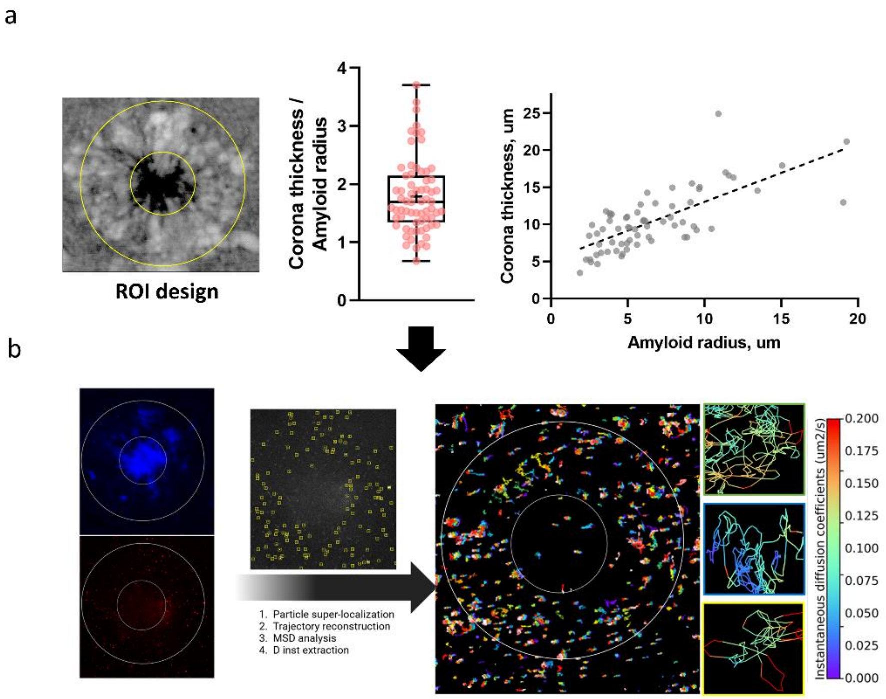


**Supplementary Figure 7.** ROI design and QDs SPT analysis (a) After conducting shadow imaging analysis, we observed that the region of interest (ROI) associated with the ring could be represented as concentric to the amyloid core, exhibiting a median width 1.82 times larger than the amyloid ratio. This relationship demonstrated a strong correlation ($r = 0.73$, $p < 0.0001$), as illustrated in Figure 2. (b) ROIs for QDs signal were drawn based on the proportions described by shadow imaging. A first ROI was drawn using the threshold blue autofluorescence signal and a second following the mentioned proportions. Analysis of QDs diffusion was performed with *PALMtracer* (MetaMorph® – Molecular Devices). Briefly, the analysis involved 1) particle super localisation, 2) trajectory reconstruction and MSD analysis, and 3) $D_{inst}$ value extraction. Each of the values was ascribed to each of the ROIs designed beforehand. The result gauges the building of a map with the trajectories of particles.

**Fig S8**

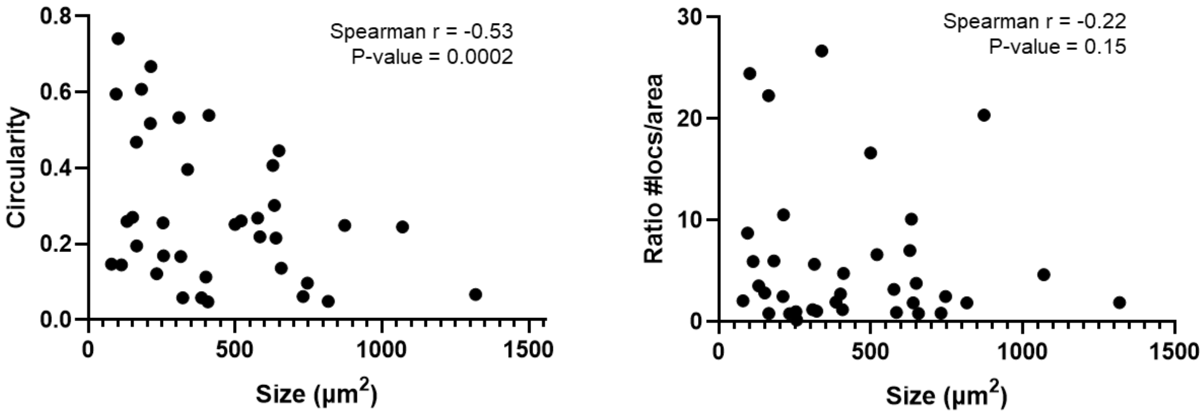


**Supplementary Figure 8.** Strong negative correlation between circularity and sizes (Spearman r =-0.53, p = 0.0002). However, there is no significant correlation between size and ratio locs/area (Spearman r = -0.22 p = 0.15).

**Fig S9**

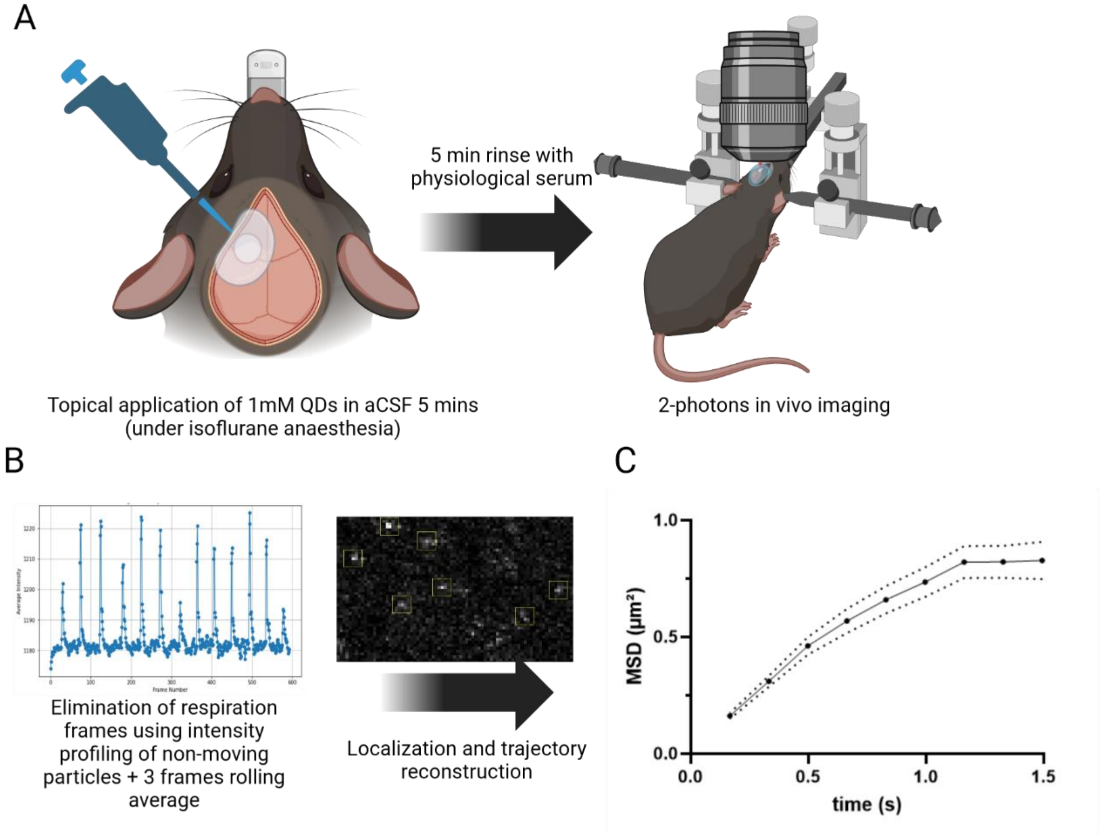


**Supplementary Figure 9. In vivo QDs SPT. (a)** A round craniotomy (1.2 mm in diameter), was made above the somatosensory cortex to expose the surface of brain. 100µL of QDs solution (1mM in aCSF) was placed over the brain surface and left for 5 minutes to diffuse into the tissue. The surface was then rinsed using physiological serum. The mice were then imaged in a 2-photon setup. (b) The analysis involved the elimination of respiration frames. A 3-point rolling average was then applied to improve signal. Localization and trajectory reconstruction were performed in MetaMorph®. (c) The total MSD shows the characteristic shape of restricted diffusion, the same found in *ex vivo*. A total of 1048 MSDs from 8 recordings (n = 2 mice) were computed in the MSD graph.

**Fig S10**

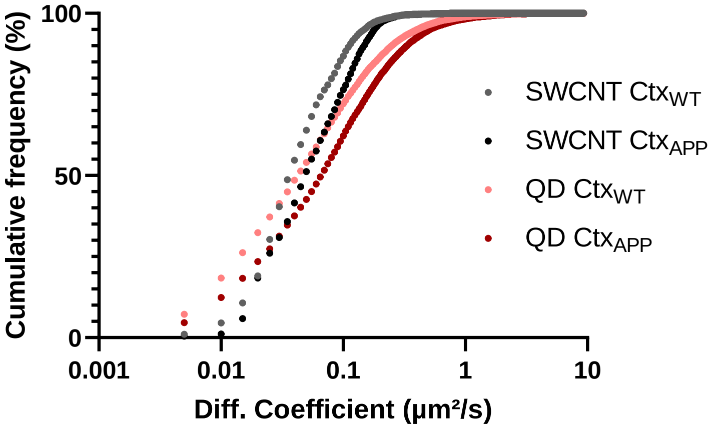


**Supplementary Figure 10.** Quantum Dots (QDs) and Single-Walled Carbon Nanotubes (SWCNTs) exhibit a comparable pattern of enhanced diffusion in the amyloid brain vs Wild-type. However, their respective instantaneous diffusion coefficients vary, likely reflecting the influence of the probe. (QDs: WT Median = 0.045, IQR = 0.017-0.115, APPPS1 Median = 0.069, IQR = 0.024-0.16; SWCNTS: WT Median = 0.038, IQR = 0.025-0.069, APPPS1 Median = 0.051, IQR = 0.027-0.98).

**Fig S11**

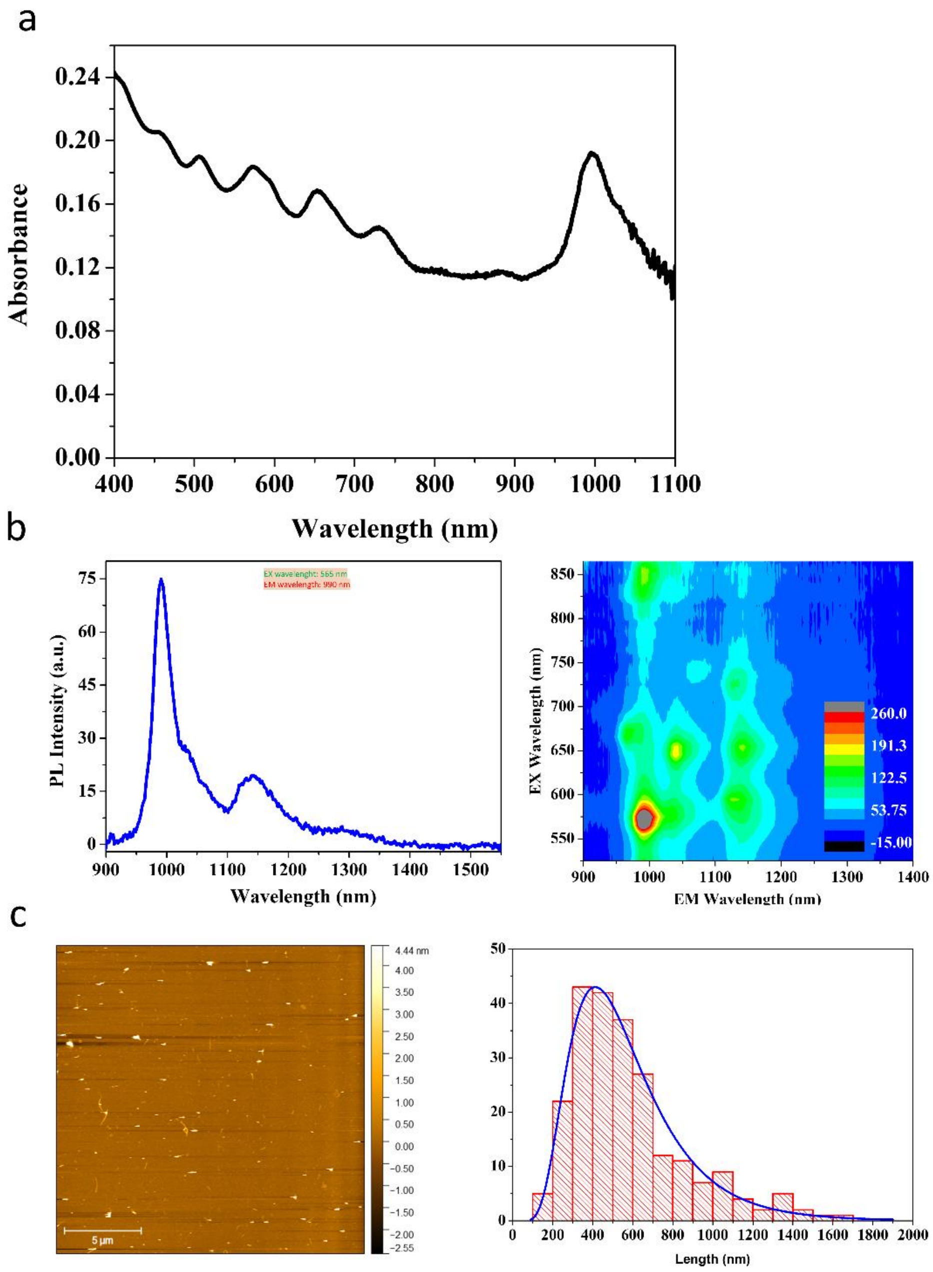


**Supplementary Figure 11.** SWCNTs spectroscopic characterisation. (a) The absorption spectrum of the HiPco synthesised CNTs suspended in 0.5% w/v PLPEG. Measured in EVOLUTION 220, Thermo Scientific (b) PL spectra of the PLPEG suspended SWCNTs with their corresponding 2D excitation-emission PL map (NanoLog, HORIBA). (c) AFM image of the PLPEG suspended SWCNTs with their length distribution. Mean length = 575 ± 285 nm (Median length = 510).

**Fig S12**

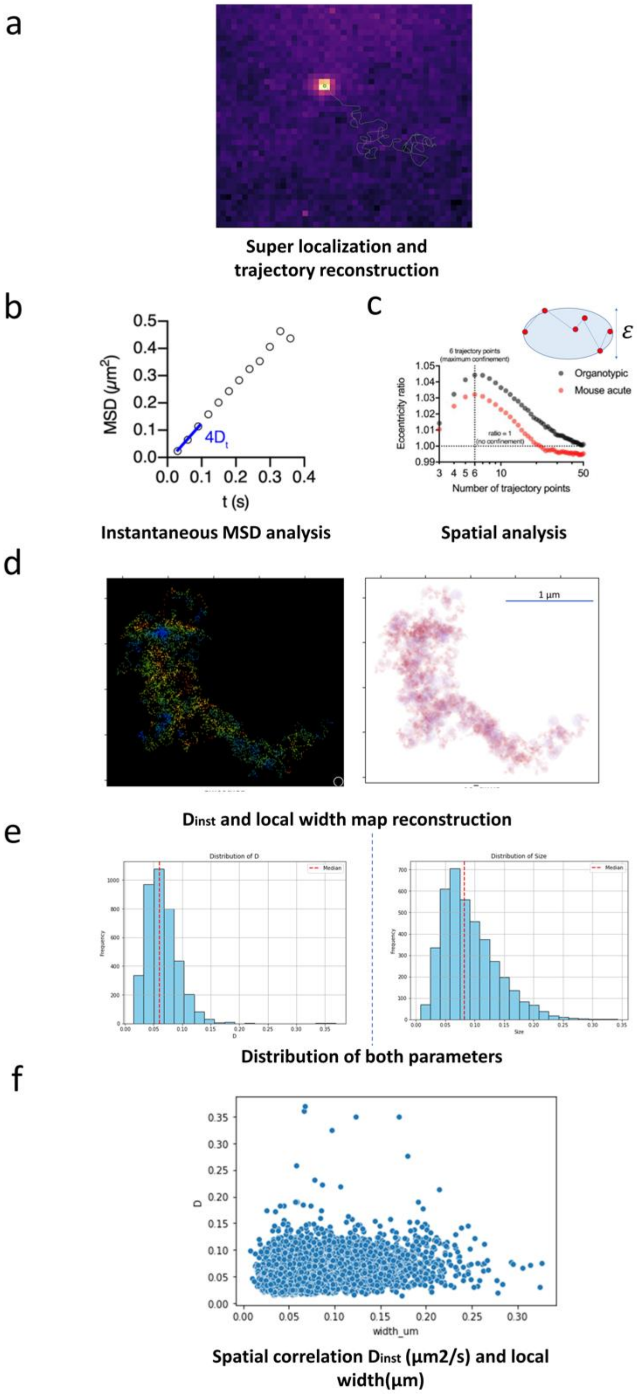


**Supplementary Figure 12**. Schematics of SWCNTs analysis. (a) Individual SWCNTs were super-localised with a 2D Gaussian fit, and coordinates were linked to reconstruct individual trajectories. (b) For each trajectory, iterative analysis of the instantaneous mean square displacement ($MSD_{inst}$) was used to estimate the instantaneous diffusion coefficient $D_{inst}$. MSD values were calculated over a sliding window of 10 frames, and linear fits were applied to the first

3 time points to retrieve $D_{inst}$'s values. (c) To estimate local ECS dimensions, the shape of the local area explored by individual SWCNTs along their trajectory was analysed using a 6-point time window. This time window was chosen at maximum confinement (i.e., when the local area's shape is maximally distorted by local ECS dimension compared to expected in unconfined environments) as defined by the eccentricity ratio of the ellipse formed by the SWCNT trajectory. Each SWCNTs trajectory rendered a (d) nanoscale map of $D_{inst}$ and local width, a (e) distribution of $D_{inst}$ and local width values and (f) spatial correlation information between both types of data.

**Fig S13**

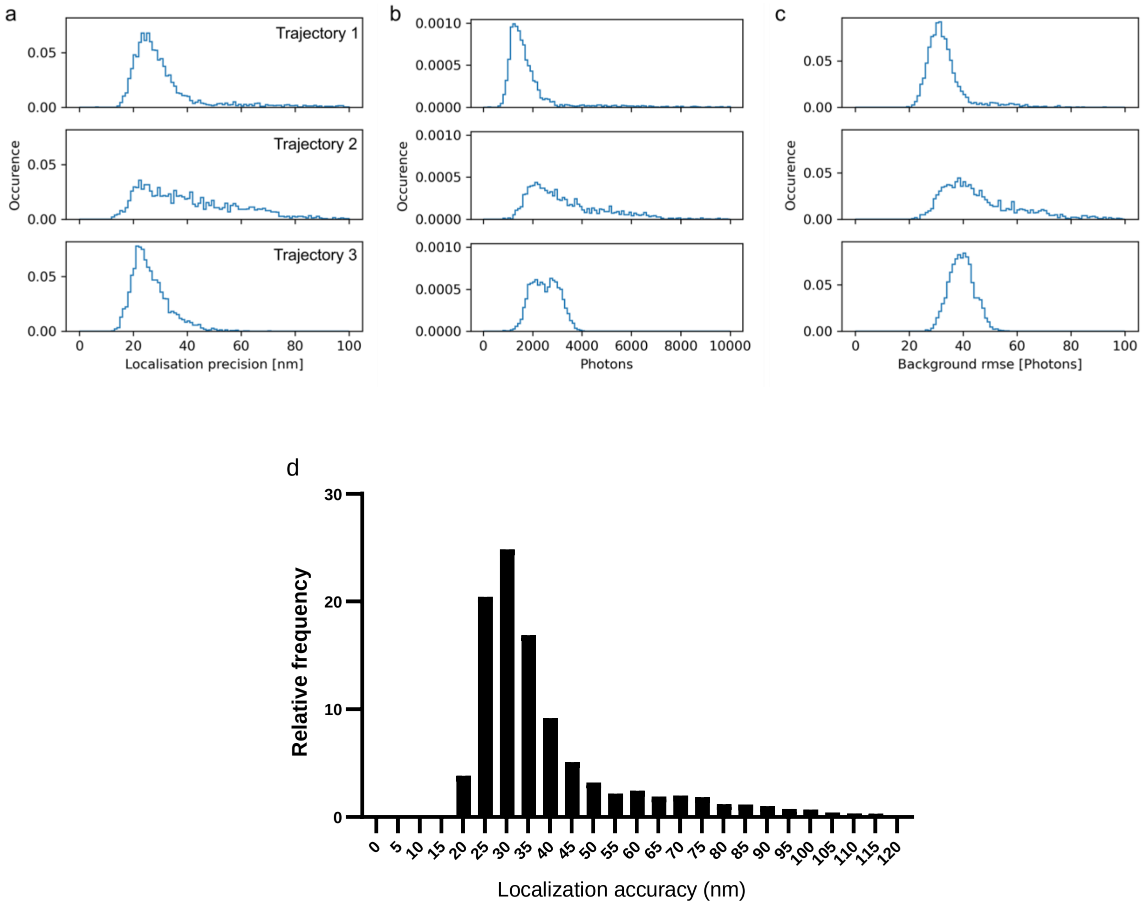


**Supplementary Figure 13**: (a) Localization precision, (b) number of photons, and (c) background noise, computed from the three representative trajectories of diffusing SWCNTs in the brain extracellular space (ECS). (d) Histogram showing localization precisions extracted from 3 previous trajecotries. Median = 32,8 IQR = 27.6-42.5.

**Supplementary Table 1. qPCR primer sequences**

| | GenBank ID | Forward Sequence (5'-3') | Reverse Sequence (5'-3') |
|---|---|---|---|
| Sdha | NM_023281 | TACAAAGTGCGGGTCGATGA | TGTTCCCCAAACGGCTTCT |
| Eef1a1 | NM_010106 | TGAACCATCCAGGCCAAATC | GCATGCTATGTGGGCTGTGT |
| Has1 | NM_008215 | AGGGCTCTTAAAGGAGGAGTCC | AGAAGGTAAACTGAGTCCCCAGAA |
| Has2 | NM_008216 | CAAAGAGGTTCGTTCAAGTTCTGA | TGTGTTTGTTTCCCACTAGCTCTC |
| Has3 | NM_008217 | CTGGTCTATCTCCTCCAACAGCTT | GCTGGGATAATGAAGAGCTACAGAA |
| Hyal1 | NM_008317 | GTGCCAAGCCCTATGCTAATAAG | GCATGTCCATTGCAAAGACTGA |
| Hyal2 | NM_010489 | GTCCCACATACACCCGAGGA | GGCACTCTCACCGATGGTAGA |
| Hyal3 | NM_178020 | GGACGACCTGATGCAGACTATTG | GGTCCCCCCAGAGTACCACT |
| Cspg4 | NM_139001 | TGGCCATGCTCTCCTCAAG | CATTCGCCACGGCTCAGT |
| Hpse | NM_152803 | AAGTTATCTTCGCAAGCCTTT | AAATGACCACAGACCTATTCCT |
| Acan | NM_001361500 | ACAGTACAATGGTTGGTACTT | CTGTCTTCTTTCAGCTCCTC |
| Vcan | NM_001081249 | CTCATCCATTCCCTTCATTTACA | ATTTCAAAGCTGGTTTCTGTTCA |
| Ncan | NM_007789 | ATCCGTAAAGGTCCCTGTG | GGTGTGGTTCCTTCTGAATTA |
| Bcan | NM_007529 | ACAAGGGCCTCAGGTTT | AAGGAGGCAGCTGACAT |
| Hapln2 | NM_022031 | TATGTCTAACCAGTCACTCAAAG | CAGACAGCTCTGGAAGTTTG |
| Hapln4 | NM_177900 | GGCTCCAGTCACACCTT | ATCCTAGCCCAGAGAAGAGT |
| Galns | NM_016722 | GGAAGATGACAGCCTAAGTTT | GAGGTGGCAGGTTACAGA |
| Mmp2 | NM_008610 | TTCCACTCCACTGCATTTCCT | GGCGGGCTGAGATGCAT |
| Mmp9 | NM_013599 | TTGCCCCTACTGGAAGGTATTATGT | GAGTGGATAGCTCGGTGGTGTT |
| Mmp16 | NM_019724 | ATAGTCTCCTGCACATCGTT | TGTCCCAACAGTCTAGCAT |
| Timp1 | NM_011593 | TGCACAGTGTTTCCCTGTTTATCTA | CCTGATCCGTCCACAAACAGT |

**Movies.**

**Supplementary video 1.** *Ex vivo* (acute slice) imaging of amyloid plaques using TUSHI in the cortex of an APP/PS1 mouse.

**Supplementary video 2.** *In vivo* imaging of amyloid plaques and ring using TUSHI in the cortex of an APP/PS1 mouse.

**Supplementary video 3.** *In vivo* imaging of amyloid plaques and ring using TUSHI in the cortex of an APP/PS1 mouse.

**Supplementary video 4.** *In vivo* imaging of the cortex using TUSHI in a WT mouse.

**Supplementary video 5.** *In vivo* imaging of the cortex using TUSHI in a WT mouse.

**Supplementary video 6.** *Ex vivo* (acute slice) imaging of QDs diffusing around a cortical amyloid plaque in an acute brain slice of an APP/PS1 mouse (Depth = 20µM).

**Supplementary video 7**. *Ex vivo* (acute slice) imaging of QDs diffusing in the cortex of a WT mouse (Depth = 20µM).

**Supplementary video 8.** *In vivo* tracking of QDs in the cortex using 2p microscopy in a WT mouse.

**Supplementary video 9.** *In vivo* tracking of QDs in the cortex using 2p microscopy in a WT mouse after breathing frames elimination and denoise (rolling average).

Biermann, B., et al., 2014. Imaging of molecular surface dynamics in brain slices using single-particle tracking. Nature communications. 5**,** 3024.
